# *X* chromosome and autosomal recombination are differentially sensitive to disruptions in SC maintenance

**DOI:** 10.1101/277764

**Authors:** Katherine Kretovich Billmyre, Cori K. Cahoon, G. Matthew Heenan, Emily Wesley, Zulin Yu, Jay R. Unruh, Satomi Takeo, R. Scott Hawley

**Author notes:** These authors contributed equally (Katherine Kretovich Billmyre, Cori K. Cahoon). Corresponding Author: R. Scott Hawley, Stowers Institute for Medical Research, Kansas City, MO 64110.

## Abstract

The synaptonemal complex (SC) is a conserved meiotic structure that regulates the repair of double strand breaks (DSBs) into crossovers or gene conversions. The removal of any central region SC component, such as the *Drosophila melanogaster* transverse filament protein C(3)G, causes a complete loss of SC structure and crossovers. To better understand the role of the SC in meiosis, we used CRISPR/Cas9 to construct three in-frame deletions within the predicted coiled-coil region of the C(3)G protein. These three deletion mutants disrupt SC maintenance at different times during pachytene and exhibit distinct defects in key meiotic processes, allowing us to define the stages of pachytene when the SC is necessary for homolog pairing and recombination. Our studies demonstrate that the *X* chromosome and the autosomes display substantially different defects in pairing and recombination when SC structure is disrupted, suggesting that the *X* chromosome is potentially regulated differently than the autosomes.

## Introduction

Several facets of meiosis ensure the faithful inheritance of chromosomes from parents to offspring. During the creation of eggs and sperm the genome must be reduced to a haploid state containing a single set of chromosomes; the failure to properly segregate chromosomes results in chromosome missegregation, leading to gametes with an incorrect number of chromosomes. Indeed, errors in meiotic chromosome segregation are the leading cause of miscarriage and aneuploidy in humans, which can result in chromosomal disorders such as Down syndrome and Turner syndrome (reviewed in (Hassold et al., 2007)).

Proper segregation of chromosomes during meiosis relies on the formation of programmed double-strand breaks (DSBs), which are initiated when the evolutionarily conserved type II DNA topoisomerase-like protein, Spo11 (Mei-W68 in *Drosophila*), forms programmed DSBs (Keeney et al., 1997; McKim and Hayashi-Hagihara, 1998). These DSBs are then repaired as crossover or gene conversion events (Fig 1A,B). Crossovers mature into chiasmata, which physically hold homologous chromosomes together from nuclear envelope breakdown until homolog separation at anaphase I, thus ensuring proper segregation of chromosomes (Nicklas, 1974). The placement of crossover events is highly non-random and is strictly regulated by multiple processes (Hughes et al., 2018). First, crossover interference prevents two crossovers from occurring in close proximity to each other (Berchowitz and Copenhaver, 2010). Second, crossovers are excluded from the heterochromatin. Third, as a result of the centromere effect, crossing over is also reduced in those euchromatic regions that lie in proximity to the centromeres (Hughes et al., 2018). Finally, even within the medial and distal euchromatin, crossing over is substantially higher toward the middle of the chromosome arms (Szauter, 1984). These constraints do not affect the frequency or distribution of gene conversion events, which appear to be randomly distributed throughout the euchromatin (Crown et al., 2018; Miller et al., 2016, 2012). Thus, the control of crossover distribution may act at the level of DSB fate choice, rather than in determining the position of DSBs.

**Figure 1:**
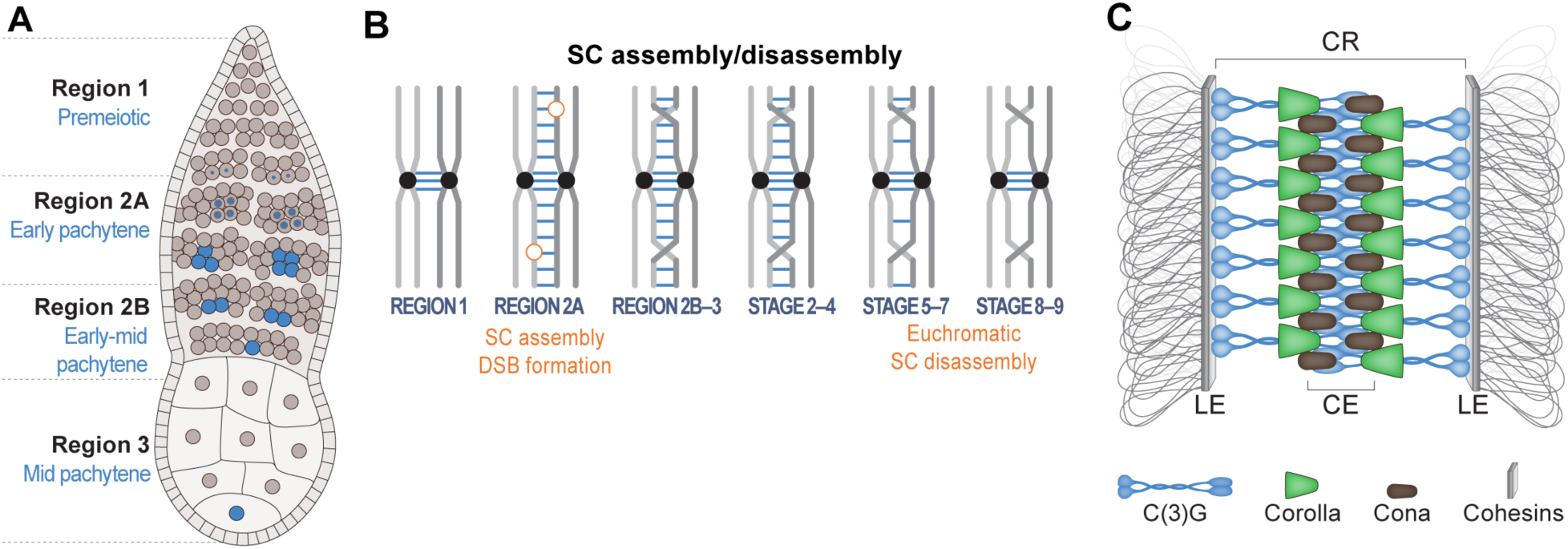
Schematic of early meiosis in *Drosophila*. (A) Diagram of a *Drosophila* germarium and SC formation (described in (Hughes et al., 2018)). At the anterior tip of the germarium, a germline stem cell divides asymmetrically to give rise to a cystoblast, which undergoes four mitotic divisions with incomplete cytokinesis to yield a 16-cell cyst. At region 2A (zygotene/early pachytene) up to 4 of the 16 cells in the cyst will enter meiosis and assemble the SC (SC represented by blue shading) to fully synapse the chromosomes. The oocyte selection process progresses in region 2B and is characterized by two nuclei (pro-oocytes) with full-length SC (early-mid pachytene) and is completed by region 3 (mid pachytene) with only one oocyte per cyst retaining full-length SC and all other nuclei having backed out of the meiotic program to become nurse cells. (B) Homologous chromosome pairing and SC assembly begin at the centromeres (represented as black dots on the chromosomes) during the mitotic divisions in region 1 (Christophorou et al., 2013; Joyce et al., 2013). In region 2A (early pachytene) the SC (represented by blue lines) is assembled along the chromosome arms and DSBs form (orange circles). The SC is maintained along chromosome arms until stage 5-7 (late pachytene), when SC disassembly occurs at multiple regions along the chromosome arms. The SC persists at the centromeres into stages 8–9 (mid prophase)(Takeo et al., 2011; Tanneti et al., 2011). (C) Model of the *Drosophila* SC showing the transverse filament protein C(3)G (blue), the central region (CR) protein Corolla (green), the central element protein CONA (black), and the lateral element/cohesin proteins (grey) connected to chromatin loops (adapted from (Hughes et al., 2018)).

Previous studies have suggested that the synaptonemal complex (SC), a large protein structure that forms between homologous chromosomes, may play a role in controlling crossover distribution (Libuda et al., 2013). The SC is a highly conserved tripartite structure, with two lateral elements and a central region (Fig 1C) (reviewed in (Cahoon and Hawley, 2016; Zickler and Kleckner, 2015, 1999)). The central region is composed of transverse filament and central element proteins, while the lateral element proteins connect the central region to the chromosome axes (Fig 1C). The known proteins that make up the *Drosophila* central region include the main transverse filament protein C(3)G, the transverse filament-like protein Corolla, and the central element protein Corona (CONA) (Collins et al., 2014; Page et al., 2008; Page and Hawley, 2001).

Work in *C. elegans* has shown that the SC functions to monitor crossover placement by preventing additional crossover designation in a region adjacent to an existing crossover precursor (Libuda et al., 2013; Nadarajan et al., 2017). Furthermore, there is evidence in *S. cerevisiae* that Zip1, a transverse filament protein, has two separable functions - one in building the SC and the other in recombination (Storlazzi et al., 1996; Voelkel-Meiman et al., 2016). Based on what is known in other model systems, it is likely that the *Drosophila* SC is also playing a role in regulating the fate of DSBs and monitoring crossover placement.

In *Drosophila*, approximately 24 DSBs are formed in early pachytene. This occurs in the context of fully formed SC after chromosome synapsis is already complete (Lake et al., 2013; Lindsley et al., 1977; Mehrotra and McKim, 2006). In the absence of the central region of the SC, DSB formation is substantially reduced, but not eliminated. Nonetheless, even in the presence of residual DSBs, there is a complete loss of crossover formation (Collins et al., 2014; Hughes et al., 2018; Mehrotra and McKim, 2006; Page et al., 2007). The abolishment of the central region of the SC also results in a high frequency of unpaired homologs during pachytene (Christophorou et al., 2013; Joyce et al., 2013; Sherizen et al., 2005; Takeo et al., 2011). In addition to disrupting meiotic pairing, the loss of any of the known central region components in the (pre-meiotic) mitotic region of the ovaries also impairs mitotic pairing of the *2nd* and *3rd* chromosomes (Christophorou et al., 2013).

Since the vast majority of SC mutants are null mutants and therefore fail to form any SC structure, it is difficult to investigate the interactions of the wildtype versions of these proteins at the protein level or discover how the SC is involved in DSB repair and fate choice. In *Drosophila*, the study of transgenes carrying in-frame deletions of either the N- or C-terminal globular domains of C(3)G have shown that both of these regions are required for proper SC assembly and crossover formation (Jeffress et al., 2007). However, these defects were too severe to allow us to investigate the function of the SC in crossover placement and formation. One domain which has not been tested is the large predicted coiled-coil domain in C(3)G. Coiled-coil domains are a key conserved feature of transverse filament proteins across many organisms and are known to be important for protein-protein interactions (Lupas and Bassler, 2017).

Here we characterize three in-frame deletion mutations in the coiled coil domain of the *Drosophila melanogaster c(3)G*, all of which cause a partial loss of SC function at different stages in early meiosis. We take advantage of the different stages of SC loss to examine when the SC is necessary for multiple meiotic events such as pairing and recombination. Unlike any previously characterized *Drosophila* meiotic mutants (Baker and Hall, 1976; Hughes et al., 2018; Parry and Sandler, 1974), the effects of these mutants on *X* chromosome recombination is different from their effects on autosomal recombination. We infer from this observation that chromosomes can respond differently to a failure in SC maintenance. We also show that the SC in early pachytene is important for the maintenance of euchromatic pairing, especially in the centromere-distal regions of the chromosome arms. The maintenance of *X* chromosome pairing is more sensitive to SC defects than is pairing maintenance on the autosomes, suggesting there may be additional chromosome-specific processes that mediate pairing. These mutants allow us for the first time to examine the temporal requirement for the synaptonemal complex in crossover placement and maintenance of pairing.

## Results

### A 213 amino acid in-frame deletion within the coiled-coil region of C(3)G impairs the maintenance of the SC in early-mid pachytene

The two previous studies of the functional anatomy of C(3)G have relied on the analysis of transgenic constructs bearing in-frame deletions (Jeffress et al., 2007; Page and Hawley, 2001). While extremely useful, transgenes have the disadvantage of non-endogenous expression levels and improper temporal expression. Based on previous studies in *S. cerevisiae* (Tung and Roeder, 1998) and in *Drosophila* (Page and Hawley, 2001), CRISPR/Cas9 was employed to construct an in-frame deletion, *c(3)G^cc^*^Δ*1*^, removing the base pairs encoding 213 amino acids (L340-A552) from the 488 amino acid predicted coil-coiled domain of C(3)G (Fig 2A, See Methods).

**Figure 2:**
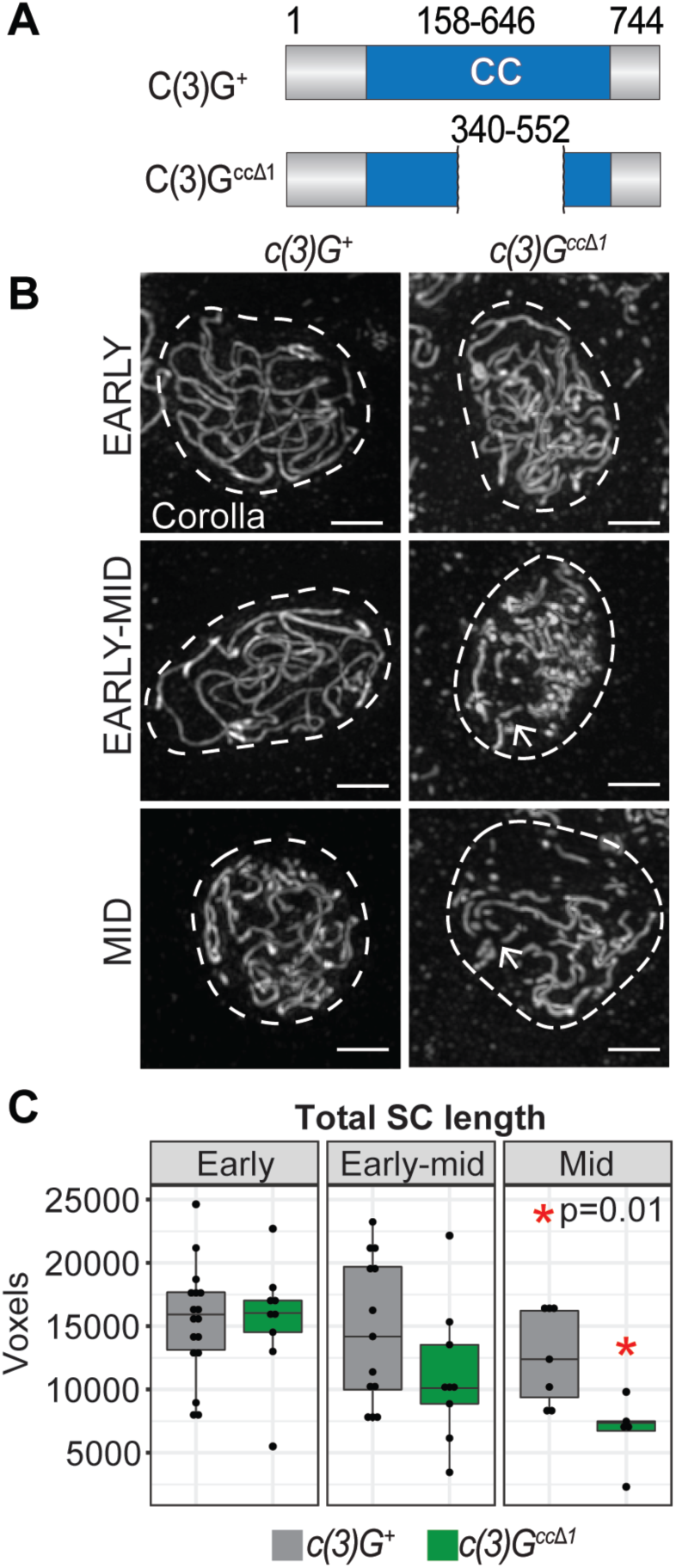
In-frame deletion of part of the large coiled-coil region of C(3)G leads to a failure to maintain SC. (A) The *c(3)G^cc^*^Δ*1*^ deletion removes the amino acids 340–552 from the coiled-coil (CC) domain of C(3)G. The predicted protein CC is in blue (based on COILs software (Lupas et al., 1991)). (B) Images showing localization of the SC protein Corolla in *c(3)G^+^* and *c(3)G^cc^*^Δ*1*^ nuclei from early pachytene (region 2A) to mid pachytene (region 3). Dotted lines indicate the location of the nucleus as defined by DAPI staining (not shown). Arrowheads indicate discontinuities in the SC. Scale bars, 2 µm. (C) Quantification of the total track length of C(3)G-positive SC in nuclei from early, early-mid and mid pachytene using skeleton analysis (See Methods). *p=0.01 by t-test, *c(3)G^+^*: N=17 (early), N=13 (early-mid), and N=7 (mid); *c(3)G^cc^*^Δ*1*^: N=9 (early), N=9 (early-mid), and N=5 (mid).

We first asked if *c(3)G^cc^*^Δ*1*^ mutants retained the ability to assemble and disassemble the SC with normal kinetics. In wild type flies, components of the central region of the SC are associated with paired centromeres during the pre-meiotic mitotic divisions (Christophorou et al., 2013; Joyce et al., 2013). By early pachytene these proteins are assembled as tripartite SC that is visible as long, continuous tracks of Corolla and C(3)G (Fig 2B, Fig 2.1A). The SC remains fully assembled until mid-late pachytene (stages 5/7), at which point the SC is removed from the euchromatic chromosome arms but remains at the centromeres in mid pachytene (Fig 1B) (reviewed (Hughes et al., 2018)). We assessed SC assembly in homozygous *c(3)G^cc^*^Δ*1*^ females using a Corolla antibody to mark the central region of the SC. In early pachytene the total length of the SC was similar to wildtype with a decrease in total SC length occurring in early-mid pachytene and a significant decrease in mid pachytene (Fig 2B,C; p=0.01). However, the SC which formed in early-mid pachytene showed obvious discontinuities (Fig 2B).

To determine whether or not the removal of a large region of the coiled-coil domain in *c(3)G^cc^*^Δ*1*^ mutants changed the tripartite structure of the SC, we measured the distance between the C-termini of C(3)G. This was accomplished using a superresolution technique, Stimulated Emission Depletion (STED), in conjunction with a C(3)G C-terminal specific antibody (Anderson et al., 2005; Collins et al., 2014). In wild type controls the distance between the C-termini of C(3)G was 118.4 nm (±0.6 nm SEM), while the distance in *c(3)G^cc^*^Δ*1*^ mutants was reduced to 67.8 nm (±0.1 nm SEM) (Fig 2.1C). The decrease in SC width is might be explained by the decreased length of C(3)G due to the 213 amino acids that were deleted. Because a single amino acid residue in a helix is predicted to be 0.15 nm in length, one would expect the decrease in length of a single C(3)G^ccΔ1^ homodimer to be 32 nm. Therefore, the width of the SC (which contains C(3)G homodimers arranged in a head to head orientation) would be predicted to be reduced by 64 nm in *c(3)G^cc^*^Δ*1*^ mutants. Although the observed 50 nm decrease in the width of the SC is less than expected, the difference may be due to differences in the way that the C(3)G^ccΔ1^ homodimer interacts with the oppositely oriented homodimer emanating from the other lateral element. Most importantly, the reduction in coiled-coil length created by removal of a large portion of the coiled-coil domain does not disrupt the formation of tripartite SC, as is illustrated by the two lateral tracks of C(3)G and the single track of Corolla observed using STED (Fig 2.1C,D).

### Loss of SC maintenance in early-mid pachytene is correlated with a reduction in *X* chromosome crossing over

The progressive (or temporal) loss of SC in *c(3)G^cc^*^Δ*1*^ flies allowed us to determine whether or not the perdurance of full-length SC until early-mid pachytene was required for proper crossing over and/or crossover placement. We examined recombination on the *X* chromosome and found that the total amount of recombination along the entire chromosome was decreased from 63 cM to 11.8 cM (Fig 3A, Table 1). This reduction in exchange was clearly polar, a well-known attribute of recombination-deficient mutants in *Drosophila* (Baker and Hall, 1976). Specifically, the chromosomal region distal and medial to the centromere from *scute (sc)* to *vermillion (v)* exhibited a very low level of crossing over (3.7% of wild type) while the centromere-proximal region from *v* to *yellow^+^ (y^+^)* was only reduced to 31.3% of wild type (Fig 3A, Table 1).

**Figure 3:**
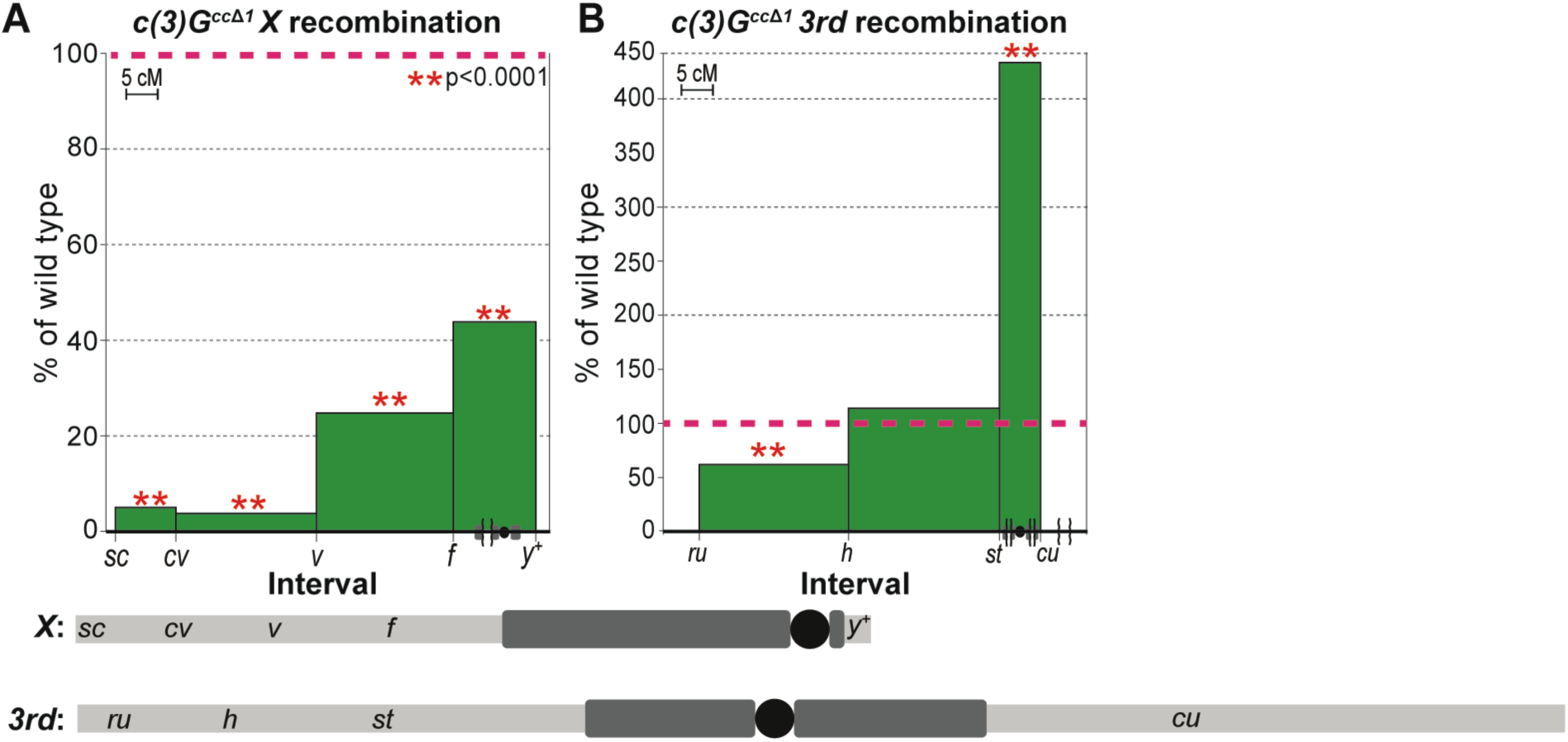
*c(3)G^cc^*^Δ*1*^ mutants exhibit chromosome-specific defects in recombination. Recombination in c*(3)G^cc^*^Δ*1*^ females on the *X* chromosome (A) and the *3rd* chromosome (B) are plotted with percent of wild type on the y-axis vs chromosome location (in cM) on the x-axis. Brackets along x-axis indicate truncation of that region of the chromosome. The red dotted line marks wild type levels of recombination and is set at 100%. P-values obtained using a Fisher’s exact test (see Table 1,2 for N values). See Methods for the recessive markers used to assay recombination. For reference, below each chart is a diagram of the corresponding chromosome being analyzed displaying the relative cytological positions of the recombination markers and the approximate amounts of pericentromeric heterochromatin estimated from (Ashburner et al., 2005) (the black circle represents the centromere).

**Table 1.** *X* Chromosome Recombination.

| Maternal genotype | $c(3)G^+$<br>(N=1515) | $c(3)G^{cc\Delta 1}$<br>(N=1420) | $c(3)G^+$<br>(N = 1721) | $c(3)G^{cc\Delta 2}$<br>(N = 1119) | $c(3)G^{cc\Delta 3}$<br>(N = 401) |
| --- | --- | --- | --- | --- | --- |
| Map Length<br>(% compared to $c(3)G^+$) | | | | | |
| $sc-cv$ | 8.8 | 0.4 (4.5%) | 10.6 | 16.6 (157%) | 0.3 (2.8%) |
| $cv-v$ | 20.7 | 0.7 (3.4%) | 18.1 | 18.5 (102%) | 0.0 (0.0%) |
| $v-f$ | 21.1 | 5.2 (24.6%) | 21.8 | 20.7 (95%) | 0.5 (2.3%) |
| $f-y^+$ | 12.4 | 5.4 (43.5%) | 11.2 | 11.5 (102%) | 2.0 (17.9%) |
| **Total** | **63.0** | **11.8 (18.7%)** | **61.7** | **67.4 (109%)** | **2.8 (4.5%)** |
| Interference |  |  |  |  |  |
| $sc/cv/v$ | 0.7 | n/a* | 0.5 | 0.8 | n/a* |
| $cv/v/f$ | 0.6 | n/a* | 0.4 | 0.5 | n/a* |
| Class |  |  |  |  |  |
| NCO | 688 | 1264 | 812 | 475 | 390 |
| SCO | 703 | 147 | 764 | 537 | 11 |
| DCO | 120 | 7 | 137 | 104 | 0 |
| TCO | 4 | 2 | 8 | 3 | 0 |
| Exchange rank |  |  |  |  |  |
| $E_0$ | 0.067 | 0.790 | 0.103 | 0.034 | 0.945 |
| $E_1$ | 0.627 | 0.196 | 0.597 | 0.604 | 0.054 |
| $E_2$ | 0.285 | 0.003 | 0.263 | 0.339 | 0 |
| $E_3$ | 0.021 | 0.011 | 0.037 | 0.021 | 0 |

Abbreviations: N, total number of flies scored; NCO, chromatids recovered exhibiting no crossovers; SCO, single-crossover chromatids; DCO, double-crossover chromatids; TCO, triple-crossover chromatids. \*Interference was not calculated unless there were at least 10 DCOs and was not calculated across the centromere. $c(3)G^+$ is $y w; pol$ .

**Table 2.** *3^rd^* Chromosome Recombination.

| Table 2. 3<sup>rd</sup> Chromosome Recombination |  |  |  |  |  |
| --- | --- | --- | --- | --- | --- |
| Maternal genotype | *c(3)G<sup>+</sup>*<br>(N=1014) | *c(3)G<sup>ccΔ1</sup>*<br>(N=1385) | *c(3)G<sup>+</sup>*<br>(N = 1994) | *c(3)G<sup>ccΔ2</sup>*<br>(N = 931) | *c(3)G<sup>ccΔ3</sup>*<br>(N = 615) |
| Map Length<br>(% compared to *c(3)G<sup>+</sup>*) |  |  |  |  |  |
| *ru-h* | 22.5 | 13.1 (58.2%) | 24.7 | 16.2 (65.5%) | 6.3 (25.5%) |
| *h-st* | 22.4 | 24.0 (107%) | 18.9 | 26.6 (141%) | 11.7 (61.9%) |
| *st-cu* | 6.1 | 27.3 (447%) | 6.8 | 23.6 (347%) | 27.5 (404%) |
| **Total** | **50.9** | **64.4 (126%)** | **50.4** | **66.4 (141%)** | **45.5 (90.3%)** |
| Interference |  |  |  |  |  |
| *ru/h/st* | 0.7 | 0.5 | 0.8 | 0.6 | n/a* |
| Class |  |  |  |  |  |
| NCO | 546 | 675 | 1069 | 418 | 377 |
| SCO | 421 | 535 | 842 | 411 | 199 |
| DCO | 45 | 167 | 83 | 99 | 36 |
| TCO | 2 | 8 | 0 | 3 | 3 |
| Exchange<br>rank |  |  |  |  |  |
| *E*<sub>0</sub> | 0.165 | 0.202 | 0.155 | 0.111 | 0.343 |
| *E*<sub>1</sub> | 0.665 | 0.420 | 0.678 | 0.477 | 0.442 |
| *E*<sub>2</sub> | 0.154 | 0.345 | 0.166 | 0.387 | 0.176 |
| *E*<sub>3</sub> | 0.016 | 0.032 | 0.000 | 0.026 | 0.039 |

Abbreviations: N, total number of flies scored; NCO, chromatids recovered exhibiting no crossovers; SCO, single-crossover chromatids; DCO, double-crossover chromatids; TCO, triple-crossover chromatids. \*Interference was not calculated unless there were at least 10 DCOs in those intervals and was not calculated across the centromere. *c(3)G<sup>+</sup>* is *y w; pol*.

The analysis of crossing over on the *3rd* chromosome did not reveal a reduction in total map length when comparing wild type and *c(3)G^cc^*^Δ*1*^ flies (Fig 3B, Table 2, 50.9 cM and 64.4 cM respectively). However, the pattern of exchange was again altered in a polar fashion, with a decrease in distal recombination between *roughoid (ru)* and *hairy (h)* (Fig 3B, Table 2, 58.2% of wild type) and a large increase (to 447% of wild type) in the centromere-proximal region between *scarlet (st)* and *curled (cu)* (Fig 3B, Table 2).

To ensure the *3rd* chromosome recombination phenotype was representative of both large autosomes, we examined recombination on the *2nd* chromosome. As shown in Fig 3.1 and Table 3, the effect of the *c(3)G^cc^*^Δ*1*^ deletion on *2nd* chromosome recombination mirrored that observed for the *3rd* chromosome with a decrease in distal recombination and a large increase on centromere-proximal exchange (Fig 3.1, Table 3). The greater than 300% increase in recombination across the centromere-proximal region on both the *2nd* and *3rd* chromosomes suggests that normal, full-length SC in early-mid and mid pachytene is regulating, directly or indirectly, crossover placement along the length of the chromosome.

**Table 3.** *2^nd^* Chromosome Recombination.

| **Table 3. 2<sup>nd</sup> Chromosome Recombination** |  |  |
| --- | --- | --- |
| Maternal genotype | *c(3)G<sup>+</sup>* | *c(3)G<sup>ccΔ1</sup>* |
|  | (N = 2376) | (N = 1456) |
| Map Length<br>(% compared to *c(3)G<sup>+</sup>*) |  |  |
| *net-dpp* | 5.7 | 3.4 (59.6%) |
| *dpp-dpy* | 8.0 | 5.8 (72.5%) |
| *dpy-b* | 28.5 | 26.9 (94.4%) |
| *b-pr* | 7.8 | 24.9 (319%) |
| *pr-cn* | 2.2 | 6.7 (304%) |
| **Total** | **52.2** | **67.6 (129%)** |
| *net/dpp/dpy* | 0.3 | 0.8 |
| *dpp/dpy/b* | 0.6 | 0.8 |
| *dpy/b/pr* | 0.6 | -0.2 |
| NCO | 1249 | 624 |
| SCO | 1021 | 692 |
| DCO | 98 | 128 |
| TCO | 8 | 12 |
| *E*<sub>0</sub> | 0.134 | 0.033 |
| *E*<sub>1</sub> | 0.715 | 0.648 |
| *E*<sub>2</sub> | 0.125 | 0.253 |
| *E*<sub>3</sub> | 0.027 | 0.066 |

Abbreviations: N, total number of flies scored; NCO, chromatids recovered exhibiting no crossovers; SCO, single-crossover chromatids; DCO, double-crossover chromatids; TCO, triple-crossover chromatids. \*Interference was not calculated unless there were at least 10 DCOs and was not calculated across the centromere. *c(3)G<sup>+</sup>* is *y w; pol*.

The striking difference in recombination patterns between the *X* chromosome and autosomes suggest that the *X* chromosome responds differently to aberrations in the SC in early-mid pachytene than the autosomes. Such chromosome-specific defects in recombination have not been previously documented in *Drosophila* (Hughes et al., 2018; Parry and Sandler, 1974).

### Smaller in-frame deletions within the putative coiled-coil domain also cause a loss of SC maintenance

One potentially confounding factor in the analysis of the *c(3)G^cc^*^Δ*1*^ mutants was the decrease in the width of the SC (Fig 2.1C) caused by the removal of a large region of the coiled-coil domain. The deletion of such a large region of the coiled coil could change the ability of the C(3)G protein to interact with itself and form stable SC but it might also remove sites important for interacting with other proteins. Therefore, in an attempt to separate the multiple phenotypes seen in *c(3)G^cc^*^Δ*1*^ flies, we created two smaller deletions within the larger deletion, *c(3)G^cc^*^Δ*2*^ (D346-T361) and *c(3)G^cc^*^Δ*3*^ (K465-V471) (Fig 4A). These smaller regions should not significantly affect the length of the C(3)G protein based on the small number of amino acids deleted. These sites were picked based on regions of C(3)G where the COILS score (Lupas et al., 1991) dipped suggesting a loss of coiled-coil structure (Fig 4.1A). We hypothesized these might be regions important for regulation of SC structure and/or function, independent of SC width.

**Figure 4:**
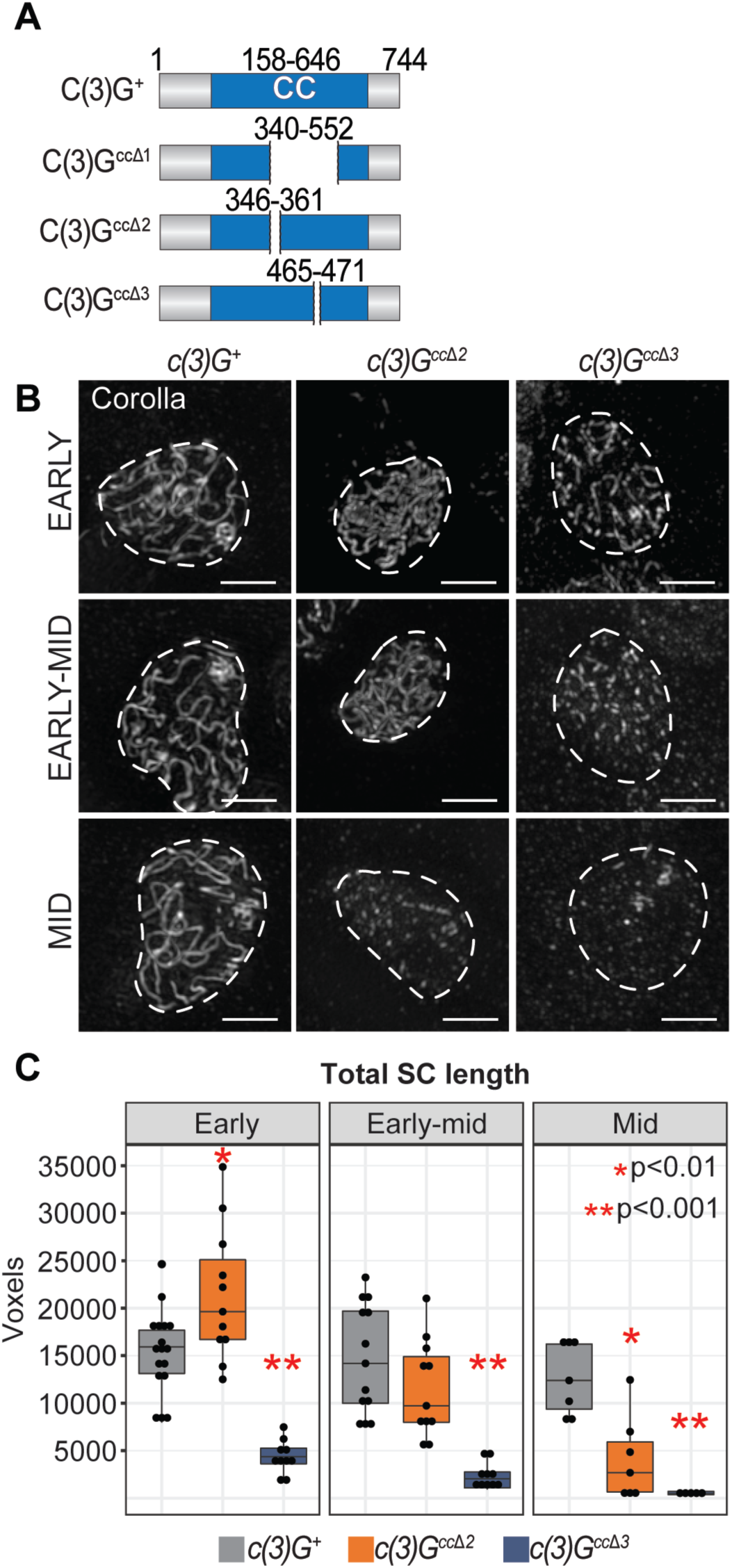
Two smaller in-frame deletions within the putative *c(3)G* coiled-coil region cause varying levels of SC defects. (A) Diagrams of the C(3)G^+^, C(3)G^ccΔ1^, C(3)G^ccΔ2^ and C(3)G^ccΔ3^ coding regions. (B) Images showing localization of the SC protein Corolla in *c(3)G^+^*, *c(3)G^cc^*^Δ*2*^, and *c(3)G^cc^*^Δ*3*^ mutants from early pachytene (region 2A) to mid pachytene (region 3). Dotted lines indicate the location of the nucleus as defined by DAPI staining (not shown). Scale bars, 2 µm. (C) Quantification of the total length of C(3)G positive SC in nuclei from early, early-mid and mid pachytene using skeleton analysis (See Methods). *c(3)G^+^* controls are the same ones used in Fig 2. *p<.01 and **p<.001 by t-test. *c(3)G^cc^*^Δ*2*^ : N=11 (early), N=11 (early-mid), and N=7 (mid); *c(3)G^cc^*^Δ*3*^: N=10 (early), N=10 (early-mid), and N=5 (mid).

When SC formation in *c(3)G^cc^*^Δ*2*^ and *c(3)G^cc^*^Δ*3*^ mutants were examined by Corolla staining, *c(3)G^cc^*^Δ*2*^ flies displayed a similar SC length to wild type in early and early-mid pachytene, but displayed a decrease in total SC length in mid pachytene when compared to wild type (Fig 4B,C, p=0.002). However, *c(3)G^cc^*^Δ*3*^ mutants never formed fully assembled full-length SC (Fig 4B,C, p<0.0001). Although each of these deletions is much smaller than the *c(3)G^cc^*^Δ*1*^ deletion, *c(3)G^cc^*^Δ*2*^ mutants did not display a loss of SC length until mid pachytene, while *c(3)G^cc^*^Δ*3*^ mutants had a more severe loss of SC in early pachytene compared to *c(3)G^cc^*^Δ*1*^ mutants (Fig 2B and 4B). We confirmed through antibody staining that the SC that did assemble in *c(3)G^cc^*^Δ*2*^ and *c(3)G^cc^*^Δ*3*^ mutants contained C(3)G (Fig 4.1B,C) in addition to Corolla (Fig 4B). The drastic differences in SC formation and maintenance observed in these mutants gave us a tool to examine the requirement of SC in early pachytene vs mid pachytene without the removal of a large structural region of C(3)G.

### Full-length SC in mid pachytene is not necessary for *X* recombination

When compared to *c(3)G^cc^*^Δ*1*^ flies, the *c(3)G^cc^*^Δ*2*^ mutants exhibited very different recombination phenotypes. First, *c(3)G^cc^*^Δ*2*^ mutants had relatively normal levels of recombination along the *X* chromosome (109% of wild type, Table 1, Fig 5A) but still displayed increased centromere-proximal recombination on the *3rd* chromosome in the *st-cu* interval (347% of wild type, Table 2,Fig 5C). Centromere-distal recombination between *ru-h* on the *3rd* chromosome was reduced to 65.5% of wild type levels in *c(3)G^cc^*^Δ*2*^ (Fig 5C, Table 2).

**Figure 5:**
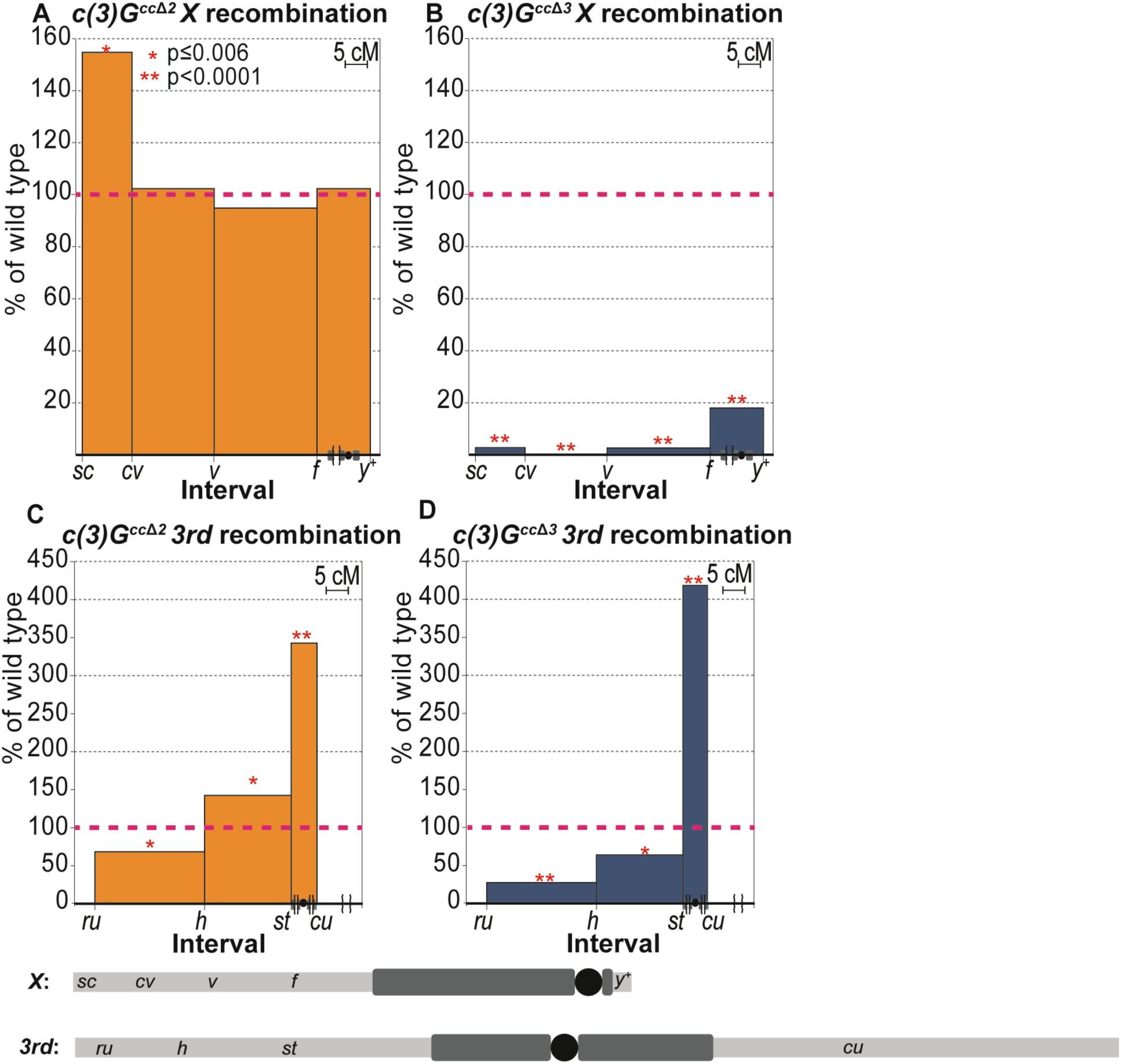
Loss of SC maintenance in *c(3)G^cc^*^Δ*2*^ mutants in mid pachytene is not sufficient to disrupt *X* chromosome recombination. Recombination in c*(3)G^cc^*^Δ*2*^ and c*(3)G^cc^*^Δ*3*^ females on the *X* chromosome (A,B) and the *3rd* chromosome (C,D) are plotted with percent of wild type on the y-axis vs chromosome location (in cM) on the x-axis. Brackets along x-axis indicate truncation of that region of the chromosome. The red dotted line marks wild type levels of recombination and is set at 100%. P-values obtained using a Fisher’s exact test (see Table 1,2 for N values). See Methods for the recessive markers used to assay recombination. For reference, below each chart is a diagram of the corresponding chromosome being analyzed displaying the relative cytological positions of the recombination markers and the approximate amounts of pericentromeric heterochromatin estimated from (Ashburner et al., 2005) (the black circle represents the centromere).

In contrast to *c(3)G^cc^*^Δ*2*^, the *c(3)G^cc^*^Δ*3*^ deletion greatly reduced recombination on the *X* chromosome to 4.5% of wild type (Fig 5B, Table 1). This reduction was similar to, but more severe than, the reduction in *X* recombination seen in *c(3)G^cc^*^Δ*1*^ mutants (18.7% of wild type, Table 1, Fig 3A). Additionally, *c(3)G^cc^*^Δ*3*^ mutants mimicked the *3rd* chromosome recombination pattern we saw in *c(3)G^cc^*^Δ*1*^ and *c(3)G^cc^*^Δ*2*^ (Fig 3B, Fig 5C, Table 2) with a centromere-distal reduction and a large centromere-proximal increase in recombination (Fig 5D, distal = 25.5% of wild type, proximal = 404% of wild type). These large increases in proximal exchange parallel those observed in *c(3)G^cc^*^Δ*1*^ mutants for both the *2nd* and *3rd* chromosomes. We note that in all cases the mutant and control crosses carry identical pericentromeric regions and therefore the observed effects on exchange in the centromere-proximal regions of the autosomes cannot be attributed to unrelated structural changes (See Methods).

### DSB formation in *c(3)G^cc^*^Δ*1*^*, c(3)G^cc^*^Δ*2*^, and *c(3)G^cc^*^Δ*3*^ mutants

To confirm that the decreases in *X* chromosome recombination observed in both the *c(3)G^cc^*^Δ*1*^ and *c(3)G^cc^*^Δ*3*^ mutants were not due to a large decrease in the formation of DSBs, we assessed DSB formation using γH2AV, a phosphorylated form of the histone variant H2AV that specifically marks sites of DSBs. Although both *c(3)G^cc^*^Δ*1*^ and *c(3)G^cc^*^Δ*3*^ flies exhibited normal kinetics for DSB repair from early to mid pachytene, *c(3)G^cc^*^Δ*1*^ flies (but not *c(3)G^cc^*^Δ*3*^ flies) displayed a decrease in the number of DSBs formed in early pachytene (Fig 5.1A, p=0.03). Because *X* chromosome recombination was more severely affected in *c(3)G^cc^*^Δ*3*^ flies compared to *c(3)G^cc^*^Δ*1*^ flies, we do not believe the early pachytene decrease in *c(3)G^cc^*^Δ*1*^ mutants is biologically relevant to the decrease in crossing over on the *X* chromosome. Lastly, we assessed DSB formation in *c(3)G^cc^*^Δ*2*^ flies and saw a slight decrease in the number of DSBs formed in early pachytene compared to wild type (Fig 5.1A, p=0.006). However, since *c(3)G^cc^*^Δ*2*^ flies did not have an overall decrease in the formation of crossovers, the decrease in γH2AV may not be biologically significant.

One possible explanation for the increase in centromere-proximal recombination might be the induction of ectopic DSBs within the heterochromatin that were not induced by Spo11. To confirm that the centromere-proximal recombination was due to Spo11 breaks, we constructed a double mutant with *c(3)G^cc^*^Δ*3*^ and *vilya^826^*, a recombination nodule component that is necessary for the induction of Spo11-induced breaks (Lake et al., 2015). When we assessed *3rd* chromosome recombination we saw very low levels of recombination (total map length = 1.4 cM, Fig 5.1B), similar to the recombination seen in *vilya^826^* alone (Lake et al., 2015). This confirmed that the crossovers in *c(3)G^cc^*^Δ*3*^ mutants are due to programmed Spo11 DSBs and not an increase in DNA damage.

### Chromosome segregation in *c(3)G^cc^*^Δ*1*^*, c(3)G^cc^*^Δ*2*^, and *c(3)G^cc^*^Δ*3*^ mutants

All previously characterized mutants in *Drosophila* that are unable to form crossovers, or have a significant reduction in crossovers genome-wide, display high levels of both *X* and *4th* chromosome nondisjunction (Collins et al., 2014; Krishnan et al., 2014; Manheim and McKim, 2003; Page and Hawley, 2001; Yan and McKee, 2013). The high levels of *X* nondisjunction observed in these recombination-defective mutants involves the interactions between both non-crossover *X* chromosomes and non-crossover autosomes (Baker and Hall, 1976; Hughes et al., 2018), such that two *X* chromosomes segregate from one autosome with the remaining autosomes segregating at random. In the absence of non-crossover autosomes, non-crossover *X* chromosomes will segregate normally.

When the rate of missegregation of the *X* and *4th* chromosomes was assessed in all three mutants, neither *c(3)G^cc^*^Δ*1*^ or *c(3)G^cc^*^Δ*2*^ mutants showed significantly increased levels of *X* or *4th* chromosome nondisjunction when compared to wild type (Table 4). *c(3)G^cc^*^Δ*3*^ mutants displayed low levels of *X* (4.5%) and *4th* (2.0%) non-disjunction (Table 4). However, this low level of non-disjunction is much lower than the 39.2% reported in *c(3)G* null mutants where the SC is completely absent (Hall, 1972).

**Table 4.** *X* and *4^th^* Chromosome Nondisjunction.

| Table 4. *X* and 4<sup>th</sup> Chromosome Nondisjunction |  |  |  |  |  |
| --- | --- | --- | --- | --- | --- |
| Maternal genotype | *c(3)G*<sup>+</sup><br>(N=1348) | *c(3)G<sup>ccΔ1</sup>*<br>(N=954) | *c(3)G*<sup>+</sup><br>(N = 1157) | *c(3)G<sup>ccΔ2</sup>*<br>(N = 2422) | *c(3)G<sup>ccΔ3</sup>*<br>(N = 837) |
| Percent nondisjunction<br>(p value) |  |  |  |  |  |
| *X* | 0.7 | 1.5 | 1.0 | 0.5 | 4.5* |
| 4<sup>th</sup> | 0.4 | 0.6 | 0.1 | 0.1 | 2.0* |

Rate of *X* and 4<sup>th</sup> nondisjunction in *c(3)G*<sup>+</sup>, *c(3)G<sup>ccΔ1</sup>*, *c(3)G<sup>ccΔ2</sup>*, and *c(3)G<sup>ccΔ3</sup>* females. Significance calculated as described in (Zeng et al., 2010). Adjusted n accounts for the inviable progeny class plus the scored progeny. \**c(3)G<sup>ccΔ3</sup>* females display elevated rates of both *X* (p < .001) and 4<sup>th</sup> (p<.001) nondisjunction when compared to *c(3)G*<sup>+</sup> controls; however the number of flies scored is not sufficient to determine significance.

The absence of an observed increase in *X* nondisjunction in *c(3)G^cc^*^Δ*1*^ and *c(3)G^cc^*^Δ*2*^ mutants is most likely explained by the absence of the nonexchange autosomes required to induce *X* chromosome nondisjunction. However, the low levels of *X* nondisjunction observed in *c(3)G^cc^*^Δ*3*^ mutants might also be compatible with a proposed role for C(3)G-like proteins in mediating achiasmate segregations (Gladstone et al., 2009; Previato de Almeida et al., 2019). Therefore, the severe SC fragmentation present in *c(3)G^cc^*^Δ*3*^ mutants may cause a mild segregation defect even in the presence of autosomal recombination.

### The loss of full-length SC in these mutants parallels the decrease in euchromatic homolog pairing

Homolog pairing is reduced in mutants lacking SC (Gong et al., 2005; Page et al., 2008; Sherizen et al., 2005). Thus, since our mutants exhibit SC defects in early to mid pachytene, we utilized them to investigate the importance of full-length SC in the maintenance of homolog pairing in *Drosophila*. Fluorescence In Situ Hybridization (FISH) was used to examine homologous pairing, and to mark the distal and proximal loci of the *X* chromosome.

In wild type, 90% to 100% of the *X* chromosome was paired from early to mid pachytene (Fig 6A). To determine what the baseline level of pairing is in the absence of the SC, *X* chromosome pairing was assessed in females homozygous for a null allele of *c(3)G* (*c(3)G^68^*). In this genotype the centromere-distal region of the chromosome was most affected, with an average of 37% paired between early and early-mid pachytene, while the centromere-proximal region was paired in about half the nuclei (Fig 6.1A, 51.5%).

**Figure 6:**
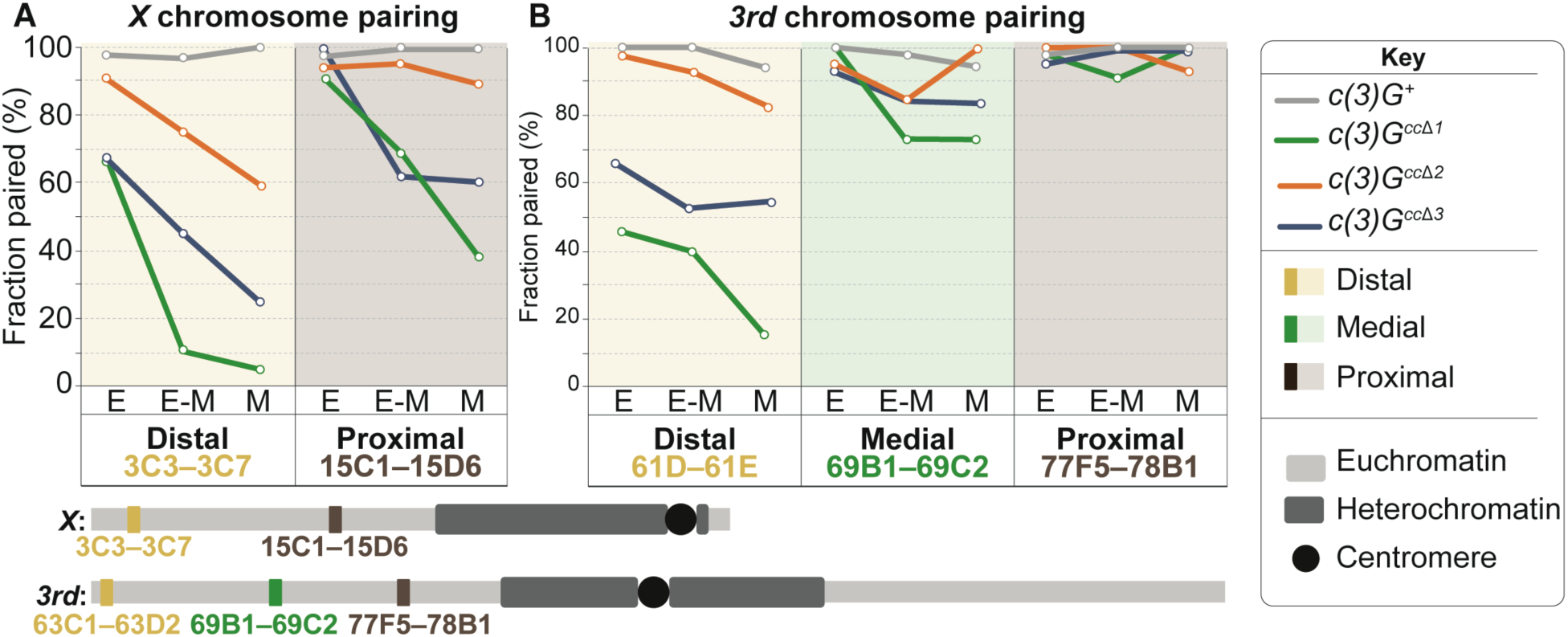
SC in early to mid pachytene maintains homologous chromosome pairing. Fraction of paired euchromatic regions in *c(3)G^+^* controls (grey line), *c(3)G^cc^*^Δ*1*^ (green line), *c(3)G^cc^*^Δ*2*^ (orange line), and *c(3)G^cc^*^Δ*3*^ flies (blue line) assessed by FISH using BAC probes against either centromere-distal or -proximal regions on the *X* chromosome (A) and centromere-distal, -medial or -proximal regions on the *3rd* chromosome (B) at early (E), early-mid (E-M) or mid (M) pachytene. For reference, below each chart is a diagram of the corresponding chromosome being analyzed (the black circle represents the centromere). For N-values see Table 5.

**Table 5.** Summary of *X* and *3^rd^* chromosome pairing.

| **Table 5. Summary of X and 3<sup>rd</sup> chromosome pairing** |  |  |  |  |  |
| --- | --- | --- | --- | --- | --- |
| Maternal genotype | *c(3)G<sup>+</sup>* | *c(3)G<sup>ccΔ1</sup>* | *c(3)G<sup>ccΔ2</sup>* | *c(3)G<sup>ccΔ3</sup>* | *c(3)G<sup>68</sup>* |
| Percent paired<br>(N value) |  |  |  |  |  |
| X proximal (3C3-3C7) |  |  |  |  |  |
| Early | 97.1(70) | 67.9(28) | 90.9(33) | 68.8(16) | n/a |
| Early-mid | 96.6(60) | 10.7(28) | 75(20) | 45(20) | 37(27) |
| Mid | 100(24) | 5.3(19) | 58.8(17) | 26.3(19) | 36.8(19) |
| X distal (15C1-15D6) |  |  |  |  |  |
| Early | 97.4(79) | 90.6(32) | 93.9(33) | 100(17) | n/a |
| Early-mid | 100(69) | 69(29) | 95(20) | 62.5(16) | 50(18) |
| Mid | 100(28) | 38.5(13) | 88.2(17) | 60(15) | 53(15) |
| 2nd proximal (22A2-22A4) |  |  |  |  |  |
| Early | 100(21) | 87.5(24) | n/a | n/a | n/a |
| Early-mid | 100(20) | 25(20) | n/a | n/a | n/a |
| Mid | 100(13) | 1(9) | n/a | n/a | n/a |
| 2nd medial (32E2-32F2) |  |  |  |  |  |
| Early | 100(21) | 89.7(29) | n/a | n/a | n/a |
| Early-mid | 100(15) | 88.2(17) | n/a | n/a | n/a |
| Mid | 100(8) | 100(7) | n/a | n/a | n/a |
| 2nd distal (38E4-38F4) |  |  |  |  |  |
| Early | 100(25) | 100(37) | n/a | n/a | n/a |
| Early-mid | 100(23) | 85.7(28) | n/a | n/a | n/a |
| Mid | 92.3(13) | 86.7(15) | n/a | n/a | n/a |
| 3rd proximal (61D-61E) |  |  |  |  |  |
| Early | 100(36) | 45.8(24) | 97.6(41) | 68.4(19) | n/a |
| Early-mid | 100(32) | 40(20) | 92.9(28) | 52.6(19) | 29.2(24) |
| Mid | 93.8(16) | 15.4(13) | 83.3(24) | 56.3(16) | 40(15) |
| 3rd medial (69B1-69C2) |  |  |  |  |  |
| Early | 100(46) | 100(33) | 95.5(22) | 93.3(30) | n/a |
| Early-mid | 97.6(43) | 73.1(26) | 85.7(14) | 86.7(15) | n/a |
| Mid | 94.4(18) | 73.6(19) | 100(9) | 84(25) | n/a |
| 3rd distal (77F5-78B1) |  |  |  |  |  |
| Early | 98.2(56) | 98.1(54) | 100(21) | 96(25) | n/a |
| Early-mid | 100(50) | 90.9(33) | 100(15) | 100(17) | 41.6(24) |
| Mid | 100(20) | 100(17) | 93.8(16) | 100(18) | 50(16) |

Starting at early-mid pachytene *c(3)G^cc^*^Δ*2*^ mutants exhibited a slight pairing defect at the centromere-distal locus of the *X* chromosome (Fig 6A, early= 90.9%, early-mid pachytene= 75%, mid pachytene= 58.8%) but were relatively well-paired at the centromere-proximal locus (Fig 6A, early=93.9%, early-mid pachytene= 95%, mid pachytene= 88.2%). Both *c(3)G^cc^*^Δ*1*^ and *c(3)G^cc^*^Δ*3*^ mutants displayed a progressive loss of pairing at both proximal and distal loci on the *X* chromosome. *c(3)G^cc^*^Δ*1*^ mutants had almost a complete loss of distal pairing by mid pachytene while *c(3)G^cc^*^Δ*3*^ mutants only maintained 26% pairing (Fig 6A).

These abnormalities in pairing maintenance correspond well with the recombination pattern seen on the *X* chromosome in *c(3)G^cc^*^Δ*1*^ and *c(3)G^cc^*^Δ*3*^ mutants in the sense that the distal region of the *X* chromosome was more affected than the proximal regions (Fig 3A and 6A). The centromere-distal decrease in recombination on the *3rd* chromosome in *c(3)G^cc^*^Δ*1*^ and *c(3)G^cc^*^Δ*3*^ mutants is displayed in conjunction with a similar loss of pairing. We examined pairing at distal, medial, and proximal loci on the *3rd* chromosome throughout pachytene. Similar to the *X* chromosome, both *c(3)G^cc^*^Δ*1*^ and *c(3)G^cc^*^Δ*3*^ mutants displayed a similar trend of reduced pairing of the *3rd* chromosome, with a progressive decrease in centromere-distal pairing that mirrors the recombination data (Fig 6B). The medial and proximal region of the *3rd* chromosome remained relatively paired from early to mid pachytene (Fig 6B). It should be noted that in *c(3)G^68^* null mutants pairing on the *3rd* chromosome was more strongly reduced; however, the proximal region (45% paired) was still paired more frequently than was the distal region (35% paired) (Fig 6.1A).

To confirm that the loss of distal pairing on the *3rd* chromosome observed in *c(3)G^cc^*^Δ*1*^ and *c(3)G^cc^*^Δ*3*^ mutants was representative of the autosomes, we also examined pairing on the *2nd* chromosome in *c(3)G^cc^*^Δ*1*^ mutants. Pairing on the *2nd* chromosome mirrored that of the *3rd* chromosome with a progressive loss of distal pairing but very little effect on medial and proximal pairing (Fig 6.1B). The significant loss of distal pairing might explain why there are stronger recombination defects in the distal regions of both the *X* and *3rd* chromosomes in *c(3)G^cc^*^Δ*1*^ and *c(3)G^cc^*^Δ*3*^ flies. By the same reason, the autosomal pairing that is maintained in these mutants is proximal, which may allow for an increased number of recombination events that are proximal to the centromere.

### Centromere pairing in meiosis is not affected in the absence of full-length SC

In wild type *Drosophila* females, the eight centromeres (two for each of the four chromosomes) pair in the pre-meiotic cysts and then cluster into an average of two clusters by early pachytene (Takeo et al., 2011). The SC is important for centromere clustering in early meiotic cells with an average of four clusters in *c(3)G, cona* and *corolla* null mutants (Collins et al., 2014; Takeo et al., 2011; Tanneti et al., 2011). Using an antibody against CID, a centromere specific histone, we assessed if centromere clustering was altered in the context of SC loss in early to mid pachytene.

Oocytes from both wild type, *c(3)G^cc^*^Δ*1*^, and *c(3)G^cc^*^Δ*2*^ flies contained an average number of clusters from 1.7 to 2.5 foci in early to mid pachytene (Fig 6.2A,B). *c(3)G^cc^*^Δ*1*^ mutants did display significantly more clusters than controls in early and mid pachytene (Fig 6.2A, p= 0.01 and 0.002 respectively). However, because the average was 2.5 foci, the loss of SC is not likely to be impacting centromere pairing in *c(3)G^cc^*^Δ*1*^. *c(3)G^cc^*^Δ*3*^ mutants had an average of 3.6 clusters in all three stages (Fig 6.2B, p< 0.001), suggesting that SC assembly defects in early pachytene may be sufficient to disrupt centromere clustering but not centromere pairing.

## Discussion

The SC plays multiple roles during meiosis that illustrate its importance in ensuring the successful transmission of genetic information from one generation to the next, yet our knowledge of how the SC is involved in regulating meiotic processes, such as recombination and the maintenance of pairing, is limited due to the integral nature of each SC component. Here we report the first partial loss-of-function SC mutations in a central region component in *Drosophila*. We use the different stages of SC loss found in these mutants to show there is a temporal requirement of the SC in the regulation of crossover number and placement on the *X* chromosome versus the autosomes (Fig 7). Additionally, full-length SC is important for maintaining euchromatic homolog pairing in distal chromosomal regions.

**Figure 7:**
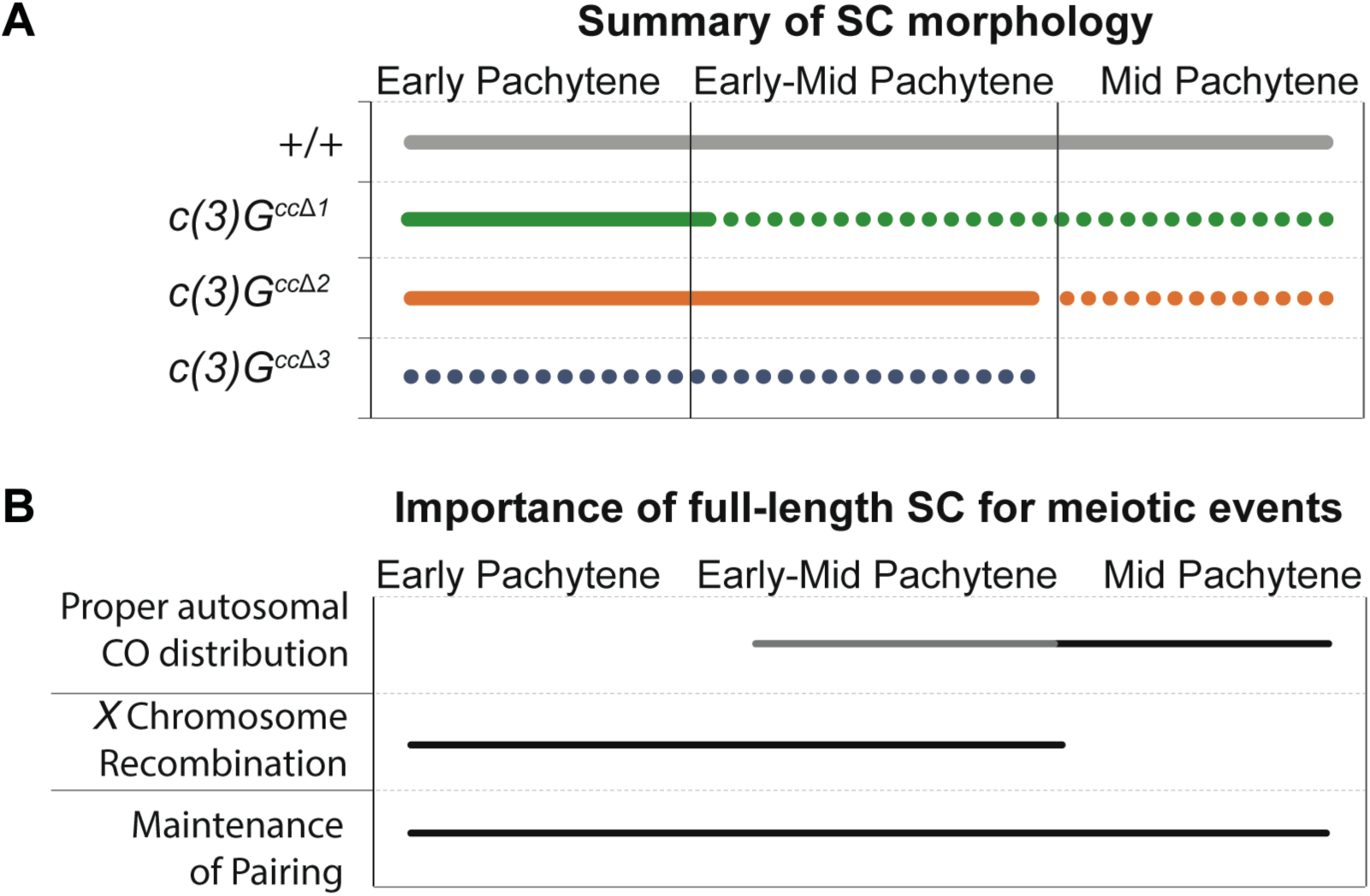
Summary of SC morphology and model of the requirement for SC in recombination and pairing maintenance. (A) Summary of SC phenotypes in *c(3)G^+^* (grey line), *c(3)G^cc^*^Δ*1*^ (green line), *c(3)G^cc^*^Δ*2*^ (orange line) and *c(3)G^cc^*^Δ*3*^ (blue line) flies. *c(3)G^cc^*^Δ*1*^ flies displayed SC defects in early-mid pachytene while *c(3)G^cc^*^Δ*2*^ flies lost SC in mid pachytene. *c(3)G^cc^*^Δ*3*^ flies never fully assembled SC. Dotted line indicates defects in total SC length and fragmentation (B) A model of the requirement of full-length SC (black lines) at different stages of pachytene. Based on our data we propose that full-length SC is important for proper autosomal crossover placement, *X* chromosome recombination and maintenance of pairing at different stages of early to mid pachytene. The grey line represents a potential role for full-length SC that cannot be confirmed with our data.

### Regulation of SC assembly and disassembly

Both the regulation of SC assembly and disassembly, and it’s maintenance after assembly, is poorly understood. Work in other organisms has shown that post-translational modifications are important in SC structure and function. It is known that SUMOylation promotes assembly of the SC while phosphorylation promotes disassembly of the SC with modifications occurring on multiple SC proteins (Jordan et al., 2012; Nadarajan et al., 2017; Sato-Carlton et al., 2017). Thus far, no post-translationally modified sites have been identified on C(3)G. However, it is likely that these sites do exist, and we speculate that sites promoting SC assembly, maintenance, and disassembly may be disrupted in these mutants.

Another possibility is that the deletions described here could destabilize protein-protein interaction sites between C(3)G and other central region proteins resulting in unstable SC that is difficult to maintain. We note that the mutant with the smallest deletion, *c(3)G^cc^*^Δ*3*^, exhibited the strongest SC defect. While this deletion was predicted to only disrupt a single coil, the best explanation for the more severe phenotype is that it actually disrupts the coiled-coil. This may have caused a large disruption in the rest of the coiled-coil structure. In the future, it will be important to further dissect these domains to better understand the regulation of SC assembly and disassembly.

### A role for the SC in the maintenance of homolog pairing

A surprising result from these studies was the ability of these deletions to allow the progressive loss of homologous euchromatic pairing through pachytene. The mechanism behind establishing and maintaining homolog pairing is a long-standing, unanswered question in the meiosis field. Previous work in *Drosophila* has shown that in the complete absence of the central region proteins C(3)G and CONA, euchromatic pairing is significantly reduced in early-mid and mid pachytene (Gong et al., 2005; Page et al., 2008; Sherizen et al., 2005).

Our partial loss-of-function mutations have allowed us to test the importance of C(3)G in maintaining pairing throughout pachytene when SC is present in early pachytene (unlike previous studies of null mutants in which the SC is always absent). From these mutants we now have a time line of when the SC is necessary to maintain pairing and recombination on the *X* chromosome and the autosomes. By comparing these mutants, we can hypothesize that the *X* chromosome needs full-length SC earlier in pachytene for proper maintenance of pairing and recombination while the autosomes are likely capable of placing crossovers as late as mid pachytene resulting in a centromere-proximal shift in crossovers where pairing is maintained (Fig 7).

In both *c(3)G^cc^*^Δ*1*^ and *c(3)G^cc^*^Δ*3*^ mutants, distal pairing of the *X* chromosome and the autosomes was most strongly reduced. One likely explanation for this is that normally the disassembly of the SC is initiated on the euchromatic chromosome arms with the centromeric region being removed last. Since the loss of the SC in c*(3)G^cc^*^Δ*1*^ and *c(3)G^cc^*^Δ*3*^ mutants occurs in a manner similar to wild type SC disassembly, the distal regions of the chromosome may be affected earlier and more strongly than the centromere-proximal regions. The centromere-proximal region contains a large amount of heterochromatin that could be mediating pairing interactions and stabilizing pairing in the absence of the SC (Dernburg et al., 1998). Furthermore, our examination of centromere pairing suggests that the centromeres are still paired (Fig 6.2) and could be facilitating the centromere-proximal pairing. This idea is supported by the higher levels of centromere-proximal pairing compared to distal pairing in *c(3)G^68^* (Fig 6.1A).

Finally, we speculate that the ability of the c*(3)G^cc^*^Δ*1*^ mutants to exhibit a centromere-distal pairing defect that is more severe than the defect seen in *c(3)G^68^* mutants results from the residual proximal crossovers that do form in *c(3)G^cc^*^Δ*1*^ mutants. Previous work has shown that crossovers can preserve synapsis but only in their vicinity (Maguire, 1985; Maguire and Riess, 1994). Perhaps the stresses that provoke separation become more concentrated on the distal regions that lack crossovers. For example, it is possible that the un-tethered distal regions could experience a higher mechanical stress due to nuclear movements than the pericentric regions containing a crossover. The lack of a strong pairing defect in c*(3)G^cc^*^Δ*2*^ mutants is probably due to the persistence of full-length SC until mid pachytene. Together these data support a role for the SC in maintaining euchromatic pairing during early to mid prophase (Fig 7).

### What causes the increase in centromere-proximal recombination events?

The autosomal increase in centromere-proximal crossovers displayed in these mutants mimics the interchromosomal effect (Crown et al., 2018; Joyce and McKim, 2009). The interchromosomal effect has been reported in flies that are heterozygous for chromosomes aberrations that suppress exchange in trans to a wild type chromosome (Lucchesi et al., 1976). Thus the absence of crossover formation on one chromosome promotes increased recombination on the other chromosomes, with more crossovers placed in the centromere-proximal regions (Crown et al., 2018; Joyce and McKim, 2009). The mechanism that controls the interchromsomal effect in balancer heterozygotes is poorly understood. It is possible that the interchromosomal effect is partially responsible for the increase in centromere-proximal crossovers in *c(3)G^cc^*^Δ*1*^ and *c(3)G^cc^*^Δ*3*^ mutants due to the loss of *X* chromosome recombination.

However, the interchromosomal effect cannot explain the increase in centromere-proximal recombination in *c(3)G^cc^*^Δ*2*^ mutants since *X* recombination appears normal. Another explanation for the increase in centromere-proximal recombination events may be the premature loss of the SC at distal regions of the chromosome. It is unknown how much of a role the SC plays in the repair of DSBs into crossover versus non-crossover events. It is possible the SC must be present to interact with factors necessary for regulating the placement of crossovers. For example, Vilya, a pro-crossover factor, localizes to the SC and DSBs prior to being recruited to recombination nodules (Lake et al., 2015). If DSB repair on the autosomes does not occur until early-mid pachytene and the SC is necessary for the determination of a crossover fate, it follows that distal loss of SC would result in a shift of crossover formation towards centromere-proximal regions where the SC is still present. This mechanism could also be increasing centromere-proximal recombination in *c(3)G^cc^*^Δ*1*^ and *c(3)G^cc^*^Δ*3*^ flies. Alternatively, SC-independent heterochromatic pairing may be holding the centromere-proximal region in close proximity allowing for crossing over in that region. In addition to interacting with pro-crossover factors the SC may be interacting with a currently unknown protein which regulates crossover placement differently on the *X* chromosome versus the autosomes.

### Why is there a difference between the *X* chromosome and the autosomes?

This set of mutants represents a unique tool to investigate not only the temporal requirements of the SC but the differences in crossover placement between the *X* chromosome and the autosomes. Since *c(3)G^cc^*^Δ*2*^ mutants do not display defects in *X* chromosome recombination we conclude that full-length SC in early-mid pachytene is necessary for *X* chromosome crossover placement (Fig 7). Examining autosomal recombination in all three mutants suggests that full-length SC is necessary in mid pachytene for proper crossover distribution on the autosomes (Fig 7). There are multiple explanations for the recombination differences between the *X* chromosome and the autosomes.

The first of these hypotheses is that there might exist a timing difference in either synapsis or crossover placement between the *X* chromosome and the autosomes. Work in *C. elegans* has provided evidence for timing differences between the sex chromosomes and the autosomes. For example, the *X* chromosome initiates pre-meiotic DNA replication later than the autosomes (Jaramillo-Lambert et al., 2007; Mlynarczyk-Evans and Villeneuve, 2017). Additionally, in *C. elegans* the *X* chromosome and the autosomes pair at the same time but synapsis of the *X* chromosome is delayed compared to the autosomes (Mlynarczyk-Evans and Villeneuve, 2017). The timing of when each chromosome is fully synapsed could be critical to ensure normal crossover placement, and the premature disruption of synapsis may affect the activity of pro-crossover factors. For example in *C. elegans*, the XND-1 protein is required for genome-wide crossover placement and is important for normal rates of DSBs on the *X* chromosome (Wagner et al., 2010). Currently, it is unknown in *Drosophila* if there are differences in the timing of DSB repair or synapsis of the *X* chromosome as compared to the autosomes, and our data suggest this as a possibility.

A second, but not mutually exclusive, explanation for the differences between the chromosomes may be a structural one. The *X* chromosome is acrocentric (centromere is near the end of the chromosome), while the autosomes are both metacentric (centromere is near the center of the chromosome), and perhaps, these structural differences mean that the *X* chromosome is more sensitive to loss of the SC. Our data suggest that loss of SC maintenance disrupts the maintenance of euchromatic homolog pairing more severely on the *X* chromosome than on the autosomes. It is unknown if metacentric chromosomes are different in terms of synapsis and recombination as compared to acrocentric chromosomes, and further investigation is needed to determine if structural differences affect these processes.

It is clear from decades of research that the regulation of recombination requires many factors and precise timing. Here we show that the SC plays a vital role in maintaining homolog pairing and proper crossover distribution in *Drosophila* female meiosis. Many differences between sex chromosomes and autosomes have been documented in a multitude of organisms, and our data are consistent with these differences extending into the processes that control chromosome pairing and recombination. With this set of mutants, we have established a new system to examine *X* chromosome and autosome biology in *Drosophila* meiosis that will allow future work to unravel the mechanism behind meiotic chromosomal differences.

## Materials and Methods

### Stocks

*Drosophila* stocks were maintained on standard food at 24°C. Descriptions of genetic markers and chromosomes can be found at http://www.flybase.org/. Wild type refers to the genotype: *y w; +/+; +/+; sv^spa-pol^*, unless stated otherwise. The key resource table contains a list of all the fly stocks used in this manuscript.

### Construction of c(3)G^ccΔ1^ mutants

To aid in screening for *c(3)G^cc^*^Δ*1*^ mutant flies, we incorporated a *piggyBac* transposon carrying a *3xP3*-DsRed that expresses in the fly eye into the intron directly downstream of the intended *c(3)G^cc^*^Δ*1*^ deletion [intron 5 of *c(3)G*] in the homologous repair template plasmid. Repair of the Cas9-induced DSB using the homologous repair plasmid will insert the desired mutation and the *piggyBac* transposon into the genome at the *c(3)G* locus. This allowed us to screen for the flies that undergo DSB repair off the homologous repair plasmid by searching for those that express dsRed in the eyes. Then, after isolation and confirmation of the desired *c(3)G* mutation, we excised the *piggyBac* transposon by crossing in a transposase. This removed any potential effect the *piggyBac* transposon may have on the expression of *c(3)G*.

The *piggyBac* transposon plasmid was constructed to have flanking *Aar*I and *Sap*I restriction sites (Addgene 51434). We used PCR to obtain two fragments of *c(3)G* from the *Drosophila* genome that flanked the position where the *piggyBac* would be inserted and added in either an *Aar*I or *Sap*I restriction site. The ~2600-bp fragment upstream of the *piggyBac* insertion site contained *Aar*I sites and was obtained using these primers: Forward, tataCACCTGCattaCCGAcgctagtggctcctagagttcag; Reverse, gcagCACCTGCgcggTTAAtgaaaaagaatttataagtcttaccattaggttatc. The ~1000-bp fragment downstream of the *piggyBac* insertion site contained *Sap*I sites and was obtained using these primers: Forward, gccgGCTCTTCNTAAccttttttctacaaaatgatttatt; Reverse, gtatGCTCTTCNCGGtcatcaaaacatagtttagtatcg.

To insert these fragments into the *piggyBac* plasmid, the plasmid and the downstream *Sap*I-containing PCR was digested with *Sap*I (also called *LguI* from ThermoFisher, ER1931), phosphatase treated (Antarctic phosphatase, NEB M0289S), and ligated together using T4 ligase (NEB, M0202S). Then, to make the *c(3)G^cc^*^Δ*1*^ mutation, the upstream *Aar*I-containing PCR fragment was TOPO cloned using the Zero Blunt TOPO kit (ThermoFisher, 451245) and cut using the restriction enzyme *HindIII* (NEB, R0104S) to remove the 702-bp fragment from *c(3)G* creating the *c(3)G^cc^*^Δ*1*^ mutation. The cut TOPO *AarI* plasmid was then phosphatase treated and ligated back together to create the *c(3)G^cc^*^Δ*1*^ deletion. Then, this plasmid was digested with AarI (ThermoFisher ER1581) to generate a ~1900-bp fragment containing the *c(3)G^cc^*^Δ*1*^ deletion, which was cloned into the *piggyBac* plasmid containing the downstream *Sap*I *c(3)G* fragment. This created the *c(3)G^cc^*^Δ*1*^ homologous repair template plasmid, which was fully sequenced to ensure all cloning occurred in the correct direction (See Key Resources for primers).

A CRISPR target sequence was selected from the flyCRISPR Optimal Target Finder (http://tools.flycrispr.molbio.wisc.edu/targetFinder/). Only a single site upstream of the *c(3)G^cc^*^Δ*1*^ deletion was selected (AAAGCTTTGTTGGCCTGTATTGG) and constructed into the pU6-BbsI-chiRNA guide RNA (gRNA) plasmid (Addgene 45946). Sense (CTTCGAAAGCTTTGTTGGCCTCTAT) and antisense (AAACATAGAGGCCAACAAAGCTTTC) oligonucleotides were ordered from IDT and cloned into the gRNA plasmid as described by the flyCRISPR subcloning pU6-gRNA protocol (http://flycrispr.molbio.wisc.edu/protocols/gRNA). After selection of the CRISPR target sequence, three single nucleotide polymorphisms (SNPs) were made in the CRISPR target sequence (the mutated bases are shown in bold: ccaataga**a**gc**g**aa**t**aaagcttt) in the *c(3)G^cc^*^Δ*1*^ homologous repair template plasmid to prevent Cas9 from cutting this plasmid. These SNPs were made using the Quik Change II XL Site-Directed Mutagenesis Kit (Agilent Technology, 200521). The gRNA and *c(3)G^cc^*^Δ*1*^ homologous repair template plasmid were sent to Genetivision (Houston, Texas) for injection into *y m*[*VASA-Cas9-3xGFP*]*ZH-2A-3xRFP w^1118^/FM7c* flies (BLM 51323). Genetivision injected the gRNA plasmid at 250 ng/µl and the *c(3)G^cc^*^Δ*1*^ homologous repair template at 500 ng/µl.

*c(3)G^cc^*^Δ*1*^ was isolated by crossing the G0 injected flies to *y w; Pr/TM3; sv^spa-pol^*, then the F1 progeny were screened for expression of dsRed in the fly eyes. Due to the *VASA*-*Cas9* transgene also being marked with RFP, only F1 males could be screened for CRIPSR insertion using dsRed expression. 15 G0 males were recovered from the commercially injected embryos (Genetivision) and crossed to *y w; Pr/TM3; sv^spa-pol^*. PCR and Sanger sequencing were used to confirm that that male had repaired off the repair template to incorporate the *c(3)G^cc^*^Δ*1*^ deletion mutation at the genomic *c(3)G* locus (See Key Resources). This was done by using forward and reverse primers that were outside of the 1kb both up and downstream repair sequence (See Key Resources). Following removal of the *piggyBac,* we sequenced the entire *c(3)G* gene to confirm both the precise excision of the transposon and that the only lesion in the gene was the desired *c(3)G^cc^*^Δ*1*^ deletion mutation. Only one male was identified and was used to establish a stock.

### Construction of c(3)G^ccΔ2^ and c(3)G^ccΔ3^ mutants

A CRISPR target sequence was selected from the flyCRISPR Optimal Target Finder (http://tools.flycrispr.molbio.wisc.edu/targetFinder/). Two guide RNAs were created for *c(3)G^cc^*^Δ*2*^ and *c(3)G^cc^*^Δ*3*^ (guide 1 *c(3)G^cc^*^Δ*2*^: GCTCAATGCGATC**T**TC**A**AGCTG**G**, guide 2 *c(3)G^cc^*^Δ*2*^: GATTGACTGATCA**G**GC**A**AC**G**AG**G**, guide 1 *c(3)G^cc^*^Δ*3*^: GCTCTTCCTG**A**TTGCTGCG**A**TG**G**, and guide 2 *c(3)G^cc^*^Δ*3*^: TCTTGAACAAC**A**ATCTGTC**A**AG**G**) and constructed into the pU6-BbsI-chiRNA guide RNA (gRNA) plasmid (Addgene 45946). Sense and antisense oligonucleotides (guide 1 *c(3)G^cc^*^Δ*2*^: CTTCGCTCAATGCGATCTTCAAGCTGG, AAACCCAGCTTGAAGATCGCATTGAGC; guide 2 *c(3)G^cc^*^Δ*2*^: CTTCGATTGACTGATCAGGCAACGAGG, AAACCCTCGTTGCCTGATCAGTCAATC; guide 1 *c(3)G^cc^*^Δ*3*^: CTTCGCTCTTCCTGATTGCTGCGATGG, AAACTCGCAGCAATCAGGAAGAGC; guide 2 *c(3)G^cc^*^Δ*3*^: CTTCTCTTGAACAACAATCTGTCAAGG, AAACTGACAGATTGTTGTTCAAGAC) were ordered from IDT and cloned into the gRNA plasmid as described by the flyCRISPR subcloning pU6-gRNA protocol (http://flycrispr.molbio.wisc.edu/protocols/gRNA).

The homologous repair constructs were created using the NEBuilder HiFi DNA Kit (NEB, E5520S) and contained 1,000 bases upstream of the first guide RNA target Cas9 site, the *c(3)G* sequence with either 42 bp (*c(3)G^cc^*^Δ*2*^) or 21 bp (*c(3)G^cc^*^Δ*3*^) removed, and 1,000 bases downstream of the second guide RNA site. The PAM sequences in the *c(3)G* gene were mutated using the Quik Change II XL Site-Directed Mutagenesis Kit (Agilent Technology). The bases changed are in bold above. Additionally, a restriction site was engineered into the repair template, without creating coding changes, to aid in genotyping (*Spe*I for *c(3)G^cc^*^Δ*2*^ and *Nhe*I for *c(3)G^cc^*^Δ*3*^).

250 ng of each gRNA plasmid and 500 ng of the homologous repair template plasmid were injected (BestGene) into *y nosCas9* (on *X* chromosome, BDSC #54591). Potential CRISPR/Cas9 hits were screened with primers (See Key Resources), which amplify a region spanning the deletion and were digested with either *Spe*I or *Nhe*I allowing for visualization of heterozygotes. Once a CRISPR/Cas9 insertion was identified, the entire *c(3)G* gene was sequenced to ensure the repair plasmid did not insert.

### Nondisjunction and recombination assays

To assay recombination along the *X* chromosome, females of the genotypes: (1) *y^1^ sc^1^ cv^1^ v^1^ f^1^ y^+^/y w; sv^spa-pol^*; 2) *y^1^ sc^1^ cv^1^ v^1^ f^1^ y^+^/y w; ru^1^ h^1^ Diap1^1^ st^1^ cu^1^ c(3)G^cc^*^Δ*1*^ *ca^1^; sv^spa-pol^/+*; 3) *y^1^ sc^1^ cv^1^ v^1^ f^1^ y^+^/y w; ru^1^ h^1^ Diap1^1^ st^1^ cu^1^ c(3)G^cc^*^Δ*2*^ *ca^1^; sv^spa-pol^/+; 4) y^1^ sc^1^ cv^1^ v^1^ f^1^ y^+^/y w; ru^1^ h^1^ Diap1^1^ st^1^ cu^1^ c(3)G^cc^*^Δ*3*^ *ca^1^; sv^spa-pol^/+*) were crossed to *y^1^ sc^1^ cv^1^ v^1^ f^1^ car^1^/B^S^Y* males. For *X* recombination analysis, only the female progeny were analyzed for the intervals *sc*-*cv*, *cv*-*v, v-f, f-y+*.

To assay recombination along the *2nd* chromosome, females of the genotypes: 1) *y w/w; net^1^ dpp^ho^ dpy^ov1^ b^1^ pr^1^ cn^1^/+; ru^1^ h^1^ Diap1^1^ st^1^ cu^1^ c(3)G^cc^*^Δ*1*^ *ca^1^/ c(3)G^cc^*^Δ*1*^ *ca^1^; sv^spa-pol^/+*; 2) *w^+^/yw; net^1^ dpp^ho^ dpy^ov1^ b^1^ pr^1^ cn^1^/*+ were crossed to *w^+^/Y; net^1^ dpp^ho^ dpy^ov1^ b^1^ pr^1^ cn^1^* males. For *2nd* recombination analysis, only the female progeny were analyzed for the intervals *net-dpp, dpp-dpy, dpy-b, b-pr, pr-cn*.

To assay recombination frequency along the *3rd* chromosome females of the following genotypes: 1) *y w/ w^+^; ru^1^ h^1^ Diap1^1^ st^1^ cu^1^ sr^1^ e^s^ ca^1^/+*; 2) *y w/ w^+^; ru^1^ h^1^ Diap1^1^ st^1^ cu^1^ c(3)G^cc^*^Δ*1*^ *ca^1^/ c(3)G^cc^*^Δ*1*^ *ca^1^; sv^spa-pol^/+*; 3) *y w/w^+^; ru^1^ h^1^ Diap1^1^ st^1^ cu^1^ c(3)G^cc^*^Δ*2*^ *ca^1^/ c(3)G^cc^*^Δ*2*^ *ca^1^; sv^spa-pol^/+*; 4) *y w/w^+^; ru^1^ h^1^ Diap1^1^ st^1^ cu^1^ c(3)G^cc^*^Δ*3*^ *ca^1^/ c(3)G^cc^*^Δ*3*^ *ca^1^; sv^spa-pol^/+*; were crossed to *w+/Y; ru^1^ h^1^ Diap1^1^ st^1^ cu^1^ sr^1^ e^s^ ca^1^* males. For *3rd* recombination analysis, only the female progeny were analyzed for the intervals *ru-h, h-st, st-cu*.

To measure the rate of both *X* and *4^th^* chromosome nondisjunction single virgin females of the indicated genotype were mated to multiple *X^Y, In(1)EN, v f B; C(4)RM, ci ey^R^* males. Calculations to determine the percentage of *X* and *4^th^* chromosome nondisjunction were performed as previously described (Hawley et al., 1992; Zitron and Hawley, 1989).

### Immunostaining of whole-mount ovaries

Germarium preparation for whole-mount immunofluorescence was modified from the protocol described in (Lake et al., 2015), with dissections performed in PBS with 0.1% Tween (PBST). Primary antibodies used included affinity-purified rabbit anti-Corolla (1:2000), mouse anti-C(3)G 1A8-1G2, 5G4-1F1, and 1G5-2F7 (all at 1:500), rabbit anti-histone H2AVD pS137 (1:500) (Rockland Inc.), mouse anti-γH2AV (1:1000) (Iowa Hybridoma Bank), rat anti-CID (used at 1:3000; gift from Claudio Sunkel), and rat anti-CID (1:500) (Hanlon et al., 2018). All secondary antibodies were used at 1:500, and the secondary antibodies used were Alexa Fluor 488 goat anti-mouse (ThermoFisher, A11001), Alexa Fluor 555 goat anti-mouse (ThermoFisher, A21422), Alexa Fluor 647 goat anti-mouse (ThermoFisher, A21235), Alexa Fluor 488 goat anti-rabbit (ThermoFisher, A11008), Alexa Fluor 555 goat anti-rabbit (ThermoFisher, A21428), Alexa Fluor 647 goat anti-rat (ThermoFisher, A21434), and Alexa Fluor 555 goat anti-rat (ThermoFisher, A21247).

For STED imaging, samples were imaged with 100x, N.A 1.40 oil. objective on a Lieca SP8 Gated STED microscope. Alexa Flour 647 labeled secondary was imaged with a pulsed white light (80 MHz) tuned to 647 nm; Alexa Fluor 594 labelled secondary was imaged with the same white laser tuned at 594 nm. Both secondaries were depleted with a pulsed STED 775 nm laser with 80-90% maximum power. All images were acquired in 2D mode to improve lateral resolution, and each image was averaged 8 times in line average mode. The emission photons were collected with an internal Leica HyD hybrid detector with a time gate between 1-6 ns. Raw STED images were deconvolved with the STED module in Huygens professional deconvolution software (version 14.10; Scientific Volume Imaging). A theoretical estimated point spread function was calculated from the raw images metadata. We used the default setting to process images, but the background was measured from raw data, also the signal to noise was set in the range of 15-20 depending on the signal intensity.

### Fluorescent in situ hybridization

FISH probes were designed from bacterial artificial chromosomes (BACs) obtained from the Children’s Hospital Oakland Research Institute (CHORI; http://bacpacresources.org/library.php?id=30). The following BACs were used: for *2L* RP98-28O9 (polytene band 22A2-22A4), RP98-43K24 (polytene band 32E2-32F2), RP98-7D17 (polytene band 38E4-38F4); for *3L* RP98-2N23 (polytene band 61D-61E), RP98-26C20 (polytene band 69B1-69C2), RP98-3J2 (polytene band 77F5-78B1); for the *X* RP98-3D13 (polytene band 3C3-3C7), RP98-9H1 (polytene band 15C1-15D6). To make the FISH probes, the BACs were PCR amplified using the Illustra GenomiPhi V2 DNA Amplification Kit (GE 25-6600-30). The concentration of the BAC DNA was determined using a Quibit and 10 ng BAC DNA was used per amplification reaction. The amplification reaction was performed via kit protocol. Next, the amplified BAC was restriction enzyme digested using *AluI* (NEB R137S), *HaeII* (NEB R107S), *MseI* (NEB R0525S), *RsaI* (NEB R0167S), *MboI* (NEB R0147S) and *MspI* (NEB R0106S). Following the digestion, the DNA was ethanol precipitated with glycogen (ThermoFisher, 10814010). The precipitated DNA was resuspended in the labeling buffer from the ULYSIS Nucleic Acid Labeling Kits (ThermoFisher – AF647 kit, U21660; AF546 kit, U21652). To label the DNA with AF647 or AF546, the protocol in the ULYSIS Nucleic Acid Labeling Kits was used with 10 µL of the digested BAC DNA. The unreacted dyes were removed from the labeling reaction using Centri-Sep Columns (Princeton Separation, CS-900).

FISH with immunohistochemistry was performed as previously described (Christophorou et al., 2013), using anti-mouse C(3)G 1A8-1G2, 5G4-1F1, and 1G5-2F7 (all at 1:500) and mouse anti-Orb antibodies 4H8 and 6H4 (1:20 each)(Developmental Studies Hybridoma Bank, Iowa). C(3)G staining was used to identify meiotic nuclei in early, early-mid or mid pachytene with the exception that in *c(3)G^cc^*^Δ*3*^ mutants mid pachytene oocytes were identified using Orb staining due to the lack of SC present. To measure the 3D distance between the FISH probe foci, a custom ImageJ plug-in (“3D jru v1”) was used with a slice spacing of 0.20 and pixel spacing of 0.06370 (available at http://research.stowers.org/imagejplugins). A locus was considered paired if the distance between the FISH probe foci was <0.75 µm and unpaired if the distanced between the FISH probe foci was ≥0.75 µm.

### Imaging and image analysis

Except for the STED imaging (see below), all images were acquired on an inverted DeltaVision microscopy system (GE Healthcare) with an Olympus 100x Objective (UPlanSApo 100x NA 1.40) and a high-resolution CCD camera or an Applied Precision OMX Blaze microscope (Issaquah, WA, USA) equipped with a PCO Edge sCMOS camera. Images were deconvolved (DeltaVision and OMX) and reconstruction was performed (OMX) using SoftWoRx v. 6.5 software (Applied Precision/GE Healthcare) following Applied Precision protocols. Images were cropped and brightness and contrast was slightly adjusted using ImageJ.

### Length measurements for the synaptonemal complex

These were performed utilizing custom macros in ImageJ (NIH, Bethesda, MD). C(3)G signals corresponding to roughly a single nucleus were traced approximately in 3D as follows. Firstly, structured illumination images were scaled in x and y by 4 with bilinear interpolation. Then they were blurred in x and y with a standard deviation of 8 pixels (80 nm). Next a rolling ball background with a radius of 50 pixels was subtracted. The resulting 3D images were thresholded at 25% of their maximum intensity to create a mask encompassing the synaptonemal complex fibers. Objects containing less than 500 voxels in 3D corresponded to noise in the image and were removed. Finally, the images were skeletonized in 3D using the 3D skeletonize plugin (based on (Lee et al., 1994), CVGIP: Graphical Models and Image Processing) to create single pixel traces of the SC in three dimensions. These were dilated once to close single pixel gaps and each 3D fiber volume was measured in voxel units for presentation. Single outliers were tested for and removed with the Grubbs test at a 1% confidence level. Statistical assessment of volume differences was accomplished with a two tailed T test.

### Line Profile Analysis of STED data

Following the profile averaging approach described in (Cahoon et al., 2017) we assessed the width of the SC. Briefly, single slice cross sectional intensity profiles were generated from manually drawn lines across regions of the SC that appeared to be flat in the z dimension (traveled along the selected plane for a substantial distance). We then aligned all of these profiles (as well as the Corolla signals where present) so that the midpoint between the C(3)G C-termini was at 0. Then, the profiles were averaged to create low noise average profile distributions. A standard t-test was used for statistical comparisons between the *c(3)G^cc^*^Δ*1*^ and wild type, the mean and standard error of the mean (SEM) were reported.

## Data and software availability

Primary data files for the figures in this paper are publicly accessible at www.stowers.org/research/publications/odr. For data analysis, the custom ImageJ plugins used are available at research.stowers.org/imagejplugins/zipped_plugins.html.

## Acknowledgements

We thank Claudio Sunkel for antibodies; past and present members of the Hawley lab for helpful discussion and comments on this manuscript; and Angela Miller for editorial and figure preparation assistance. R.S.H. is an American Cancer Society Research Professor.

---

**Figure 2-figure supplement 1:**
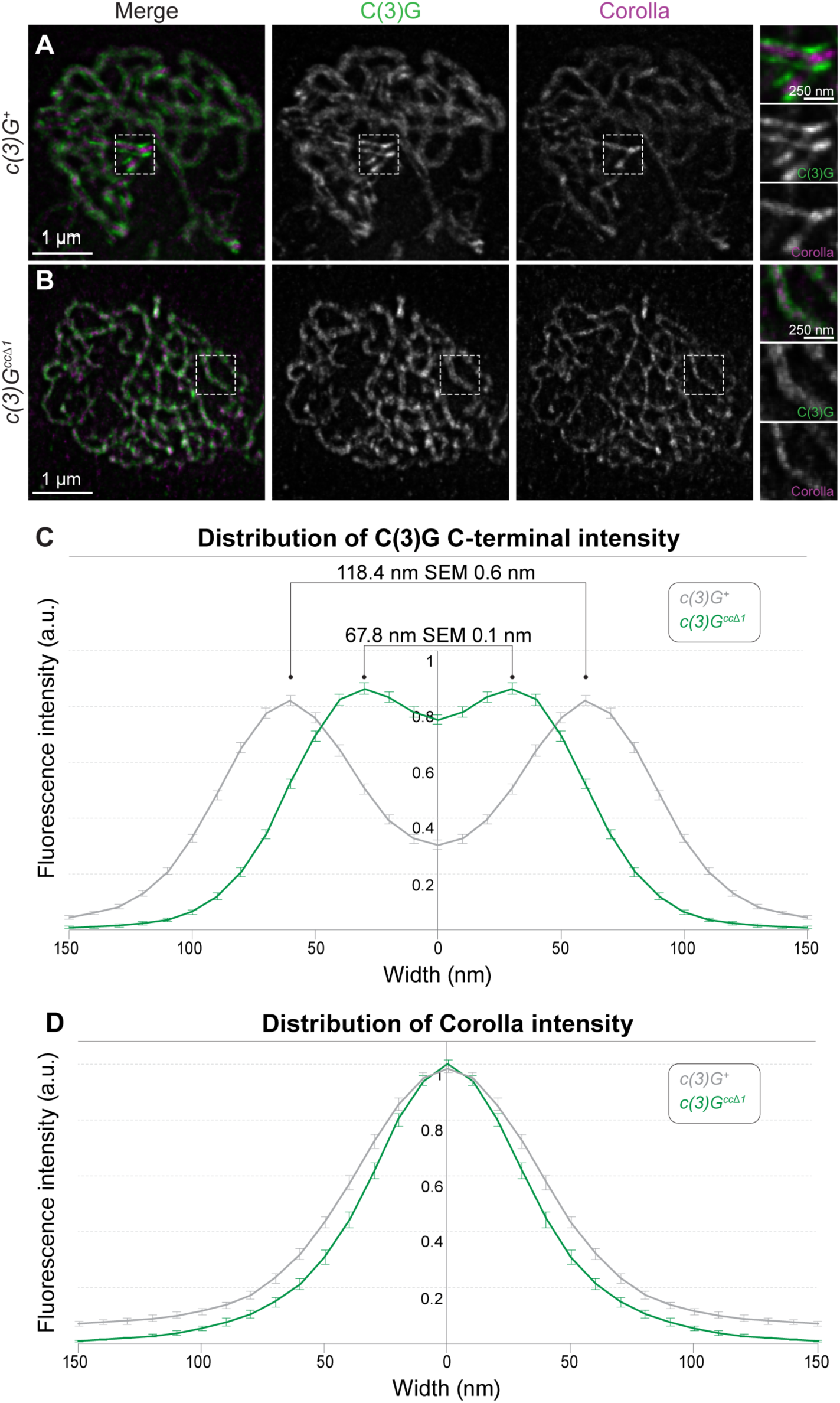
Width of SC is reduced, but the tripartite structure is maintained, in *c(3)G^cc^*^Δ^*^1^* mutants in early pachytene. STED images of early pachytene nuclei with the C-terminus of C(3)G (green) and Corolla (magenta) labeled in *c(3)G^+^* (A) and *c(3)G^cc^*^Δ*1*^ mutants (B). (C)The average distribution of the distance between the two C-terminal C(3)G tracks is shown based on a line profile analysis of STED data in each genotype (see Methods). The quantification resulted in an average width of 118.4 nm ± 0.6 nm (SEM) in wild type and 67.8 nm ± 0.1 nm (SEM) in *c(3)G^cc^*^Δ*1*^ mutants. (D) The average distribution of the Corolla signal based on a line profile analysis of STED data in each genotype. The average distribution was generated by averaging 46 line profiles from 8 wild type nuclei and 35 line profiles from 12 *c(3)G^cc^*^Δ*1*^ nuclei.

**Figure 3-figure supplement 1:**
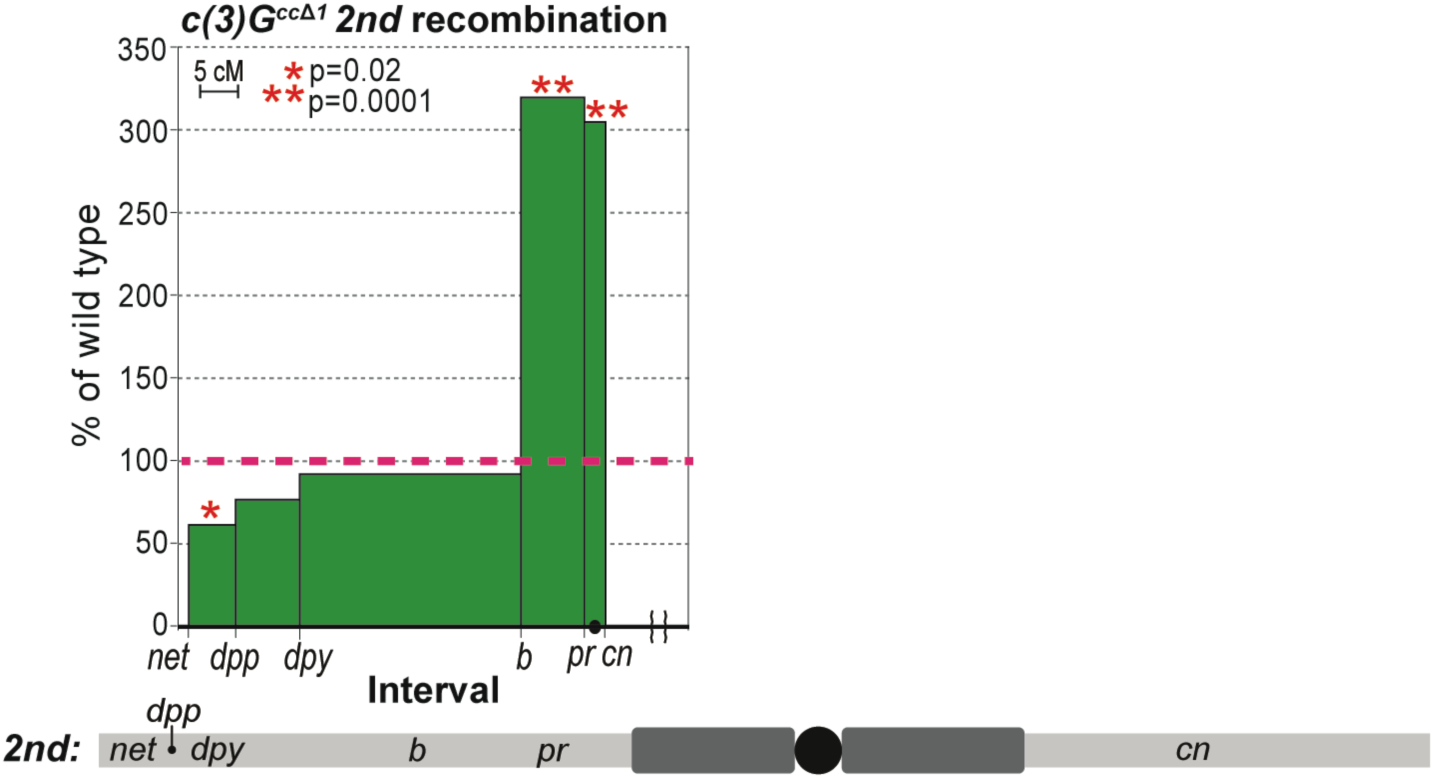
*c(3)G^cc^*^Δ*1*^ mutants exhibit recombination defects on the *2nd* chromosome. Recombination in *c(3)G^cc^*^Δ*1*^ mutants on the *2nd* chromosome plotted with percent of wild type on the y-axis vs chromosome location (in cM) on the x-axis. Brackets along x-axis indicate truncation of that region of the chromosome. The red dotted line marks wild type levels of recombination and is set to 100%. See Methods for the recessive markers used to assay recombination. P-values obtained using a Fisher’s exact test (see Table 3 for N values). For reference, below each chart is a diagram of the corresponding chromosome being analyzed displaying the relative cytological positions of the recombination markers and the approximate amounts of pericentromeric heterochromatin estimated from (Ashburner et al., 2005) (the black circle represents the centromere).

**Figure 4-figure supplement 1:**
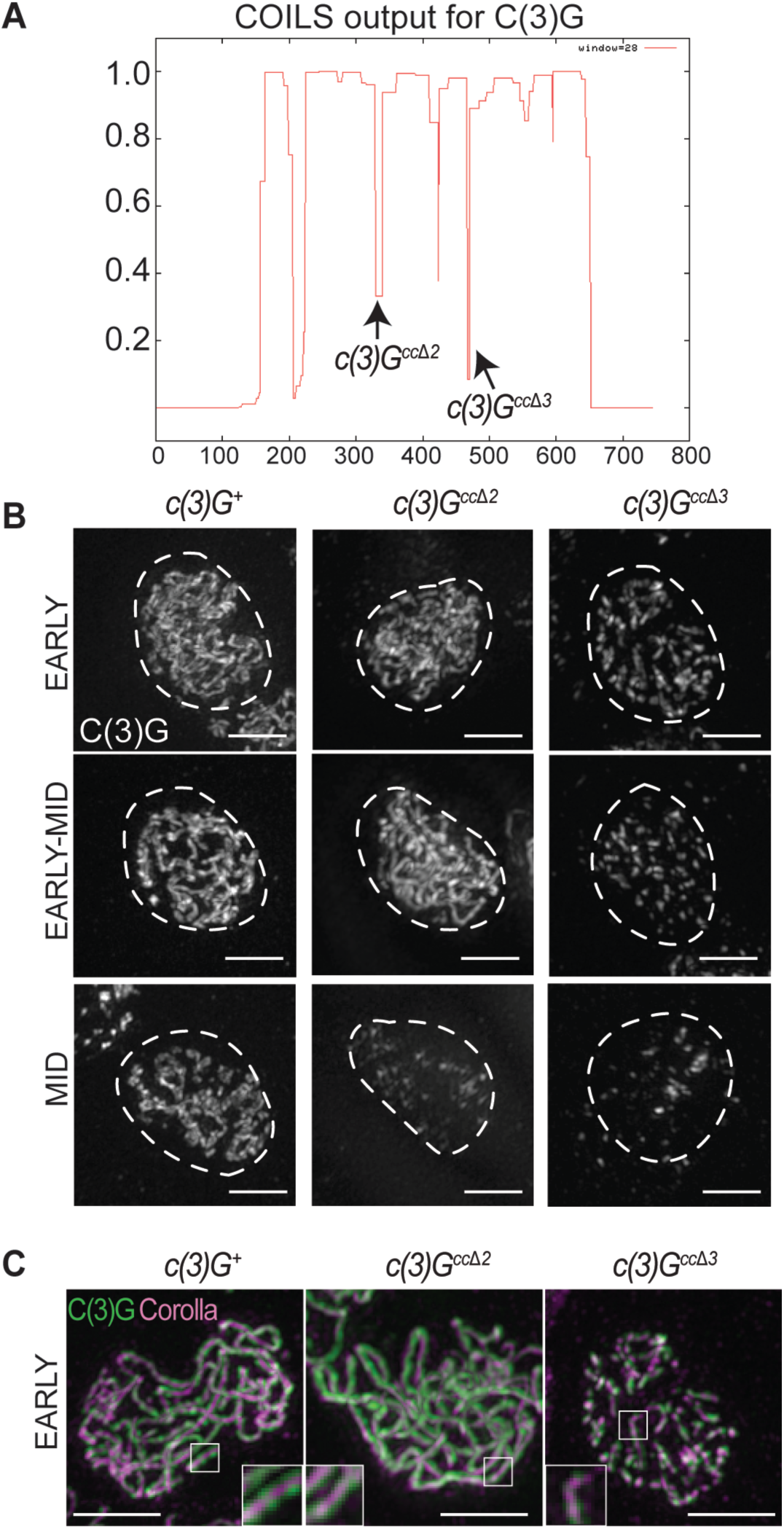
C(3)G is present in *c(3)G^cc^*^Δ*2*^ and *c(3)G^cc^*^Δ*3*^ mutants. (A) Diagram of COILS score along the length of C(3)G with arrows where *c(3)G^cc^*^Δ*2*^ and *c(3)G^cc^*^Δ*3*^ deletions were made (Lupas et al., 1991). (B) Images showing localization of the SC protein C(3)G in wild type, *c(3)G^cc^*^Δ*2*^, and *c(3)G^cc^*^Δ*3*^ flies from early pachytene (region 2A) to mid pachytene (region 3). Dotted lines indicate the location of the nucleus as defined by DAPI staining (not shown). (C) Combined Corolla (magenta) and C(3)G (green) staining in early pachytene nuclei show localization of Corolla to the middle of C(3)G. Scale bars, 2 µm.

**Figure S5-figure supplement 1:**
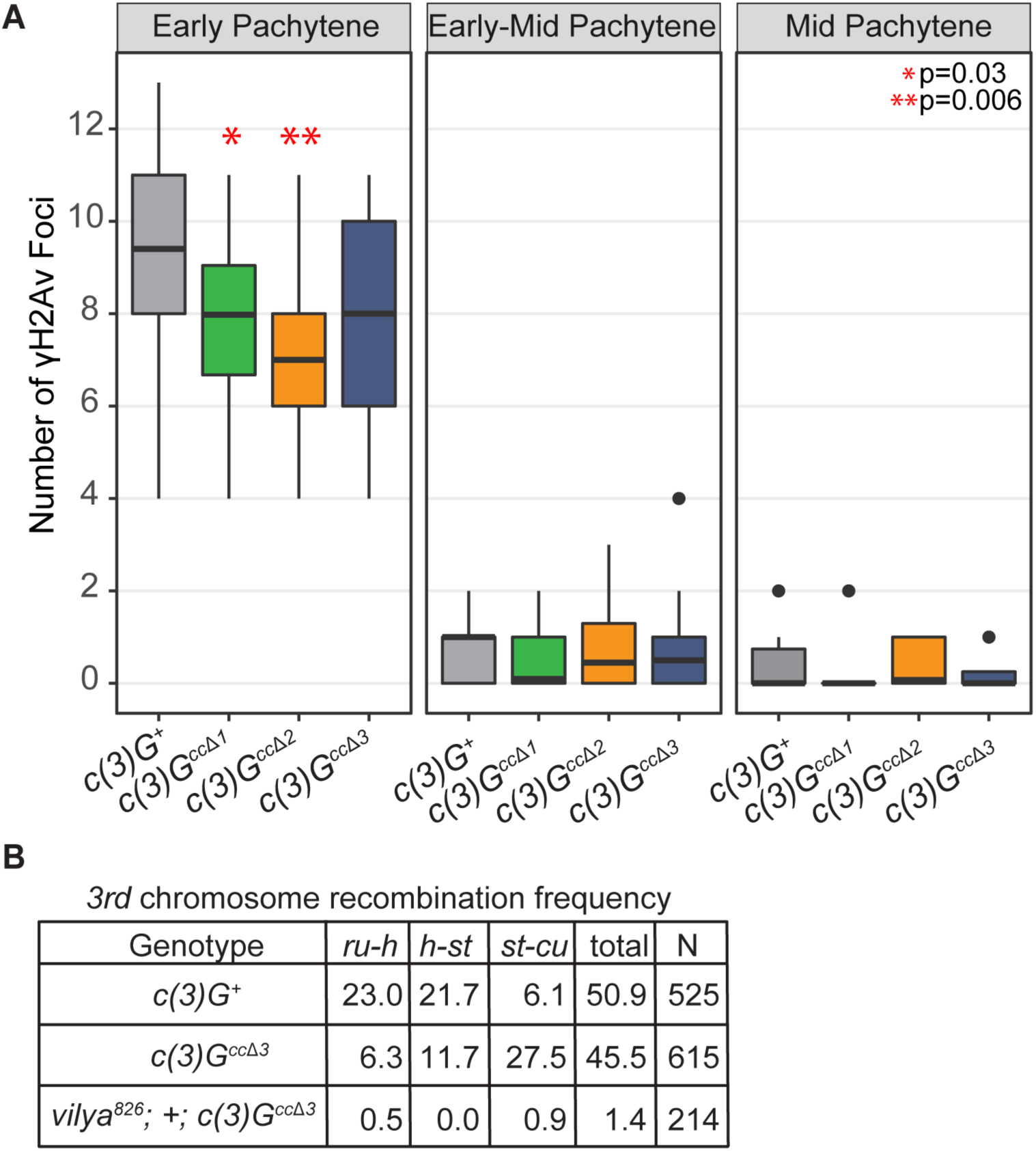
DSB levels, as determined by γH2AV foci number, in *c(3)G^cc^*^Δ*1*^, *c(3)G^cc^*^Δ*2*^ and *c(3)G^cc^*^Δ*3*^ mutants are similar to wild type. (A) Quantification of the number of DSBs per nucleus was determined by counting the number of γH2AV foci in early, early-mid and mid pachytene of the germarium for *c(3)G^+^*, *c(3)G^cc^*^Δ*1*^, *c(3)G^cc^*^Δ*2*^ and *c(3)G^cc^*^Δ*3*^ flies. N value is ≥ 10 nuclei. Statistics were performed using the Mann-Whitney test. (B) *3rd* chromosome recombination frequency in *c(3)G^+^*, *c(3)G ^cc^*^Δ*3*^ and, *vilya^826^;c(3)G ^cc^*^Δ*3*^ double mutants. Data for *c(3)G ^cc^*^Δ*3*^ is the same as shown in Figure 5 and Table 2.

**Figure S6-figure supplement 1:**
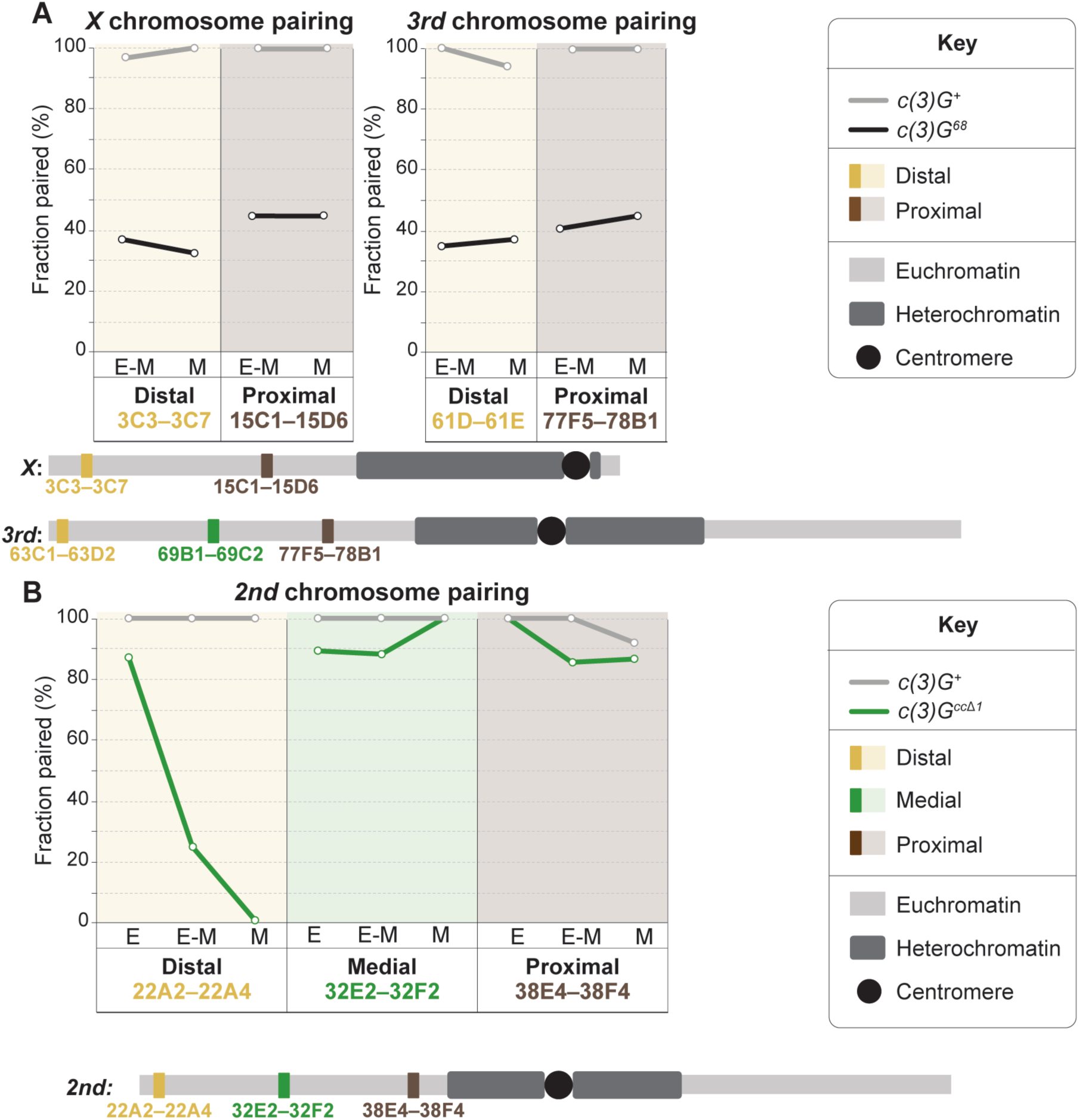
Pairing in *c(3)G^68^* and *c(3)G^cc^*^Δ*1*^ mutants. (A) Fraction of paired euchromatic regions assessed by FISH using BAC probes against centromere-distal and proximal regions of the *X* and *3rd* chromosomes in *c(3)G^68^* (black line) mutants at early-mid (E-M) or mid (M) pachytene. *c(3)G^+^* control data was previously presented in Figure 6. (B) Fraction of paired euchromatic regions on the *2nd* chromosome at early (E), early-mid (E-M) or mid (M) pachytene in *c(3)G^cc^*^Δ*1*^ (green line) mutants compared to *c(3)G^+^* (grey line) controls. See Table 5 for N values.

**Figure 6-figure supplement 2:**
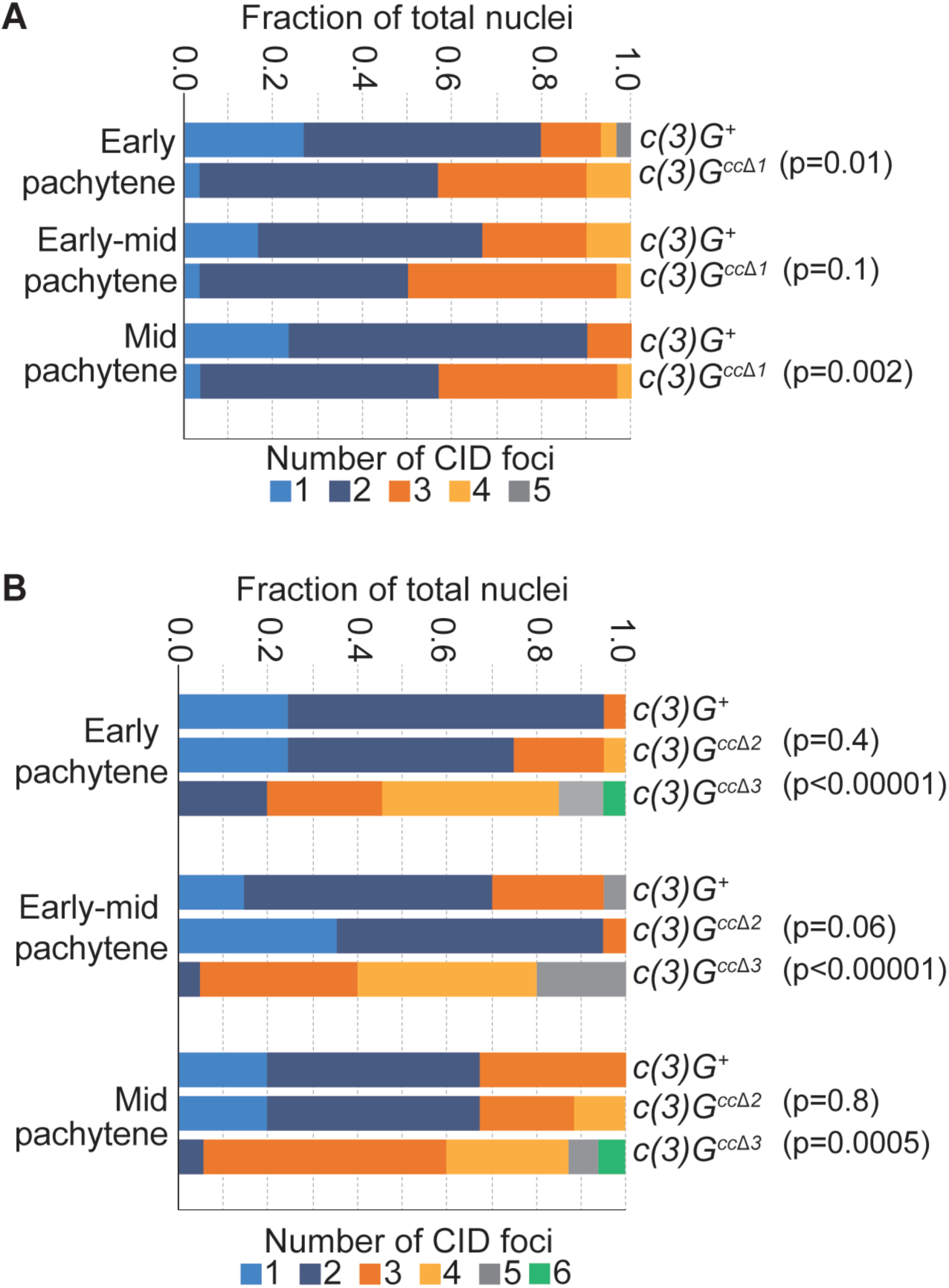
Centromere pairing *c(3)G^cc^*^Δ*1*^, *c(3)G^cc^*^Δ*2*^, and *c(3)G^cc^*^Δ*3*^ mutants. Quantification of the number of CID foci per nucleus in wild type, *c(3)G^cc^*^Δ*1*^(A), *c(3)G^cc^*^Δ*2*^ (B), and *c(3)G^cc^*^Δ*3*^ mutants (B) from early pachytene (region 2A) to mid pachytene (region 3) shows no loss of centromere pairing. Statistics were performed using the Mann-Whitney test. N ≥ 15.

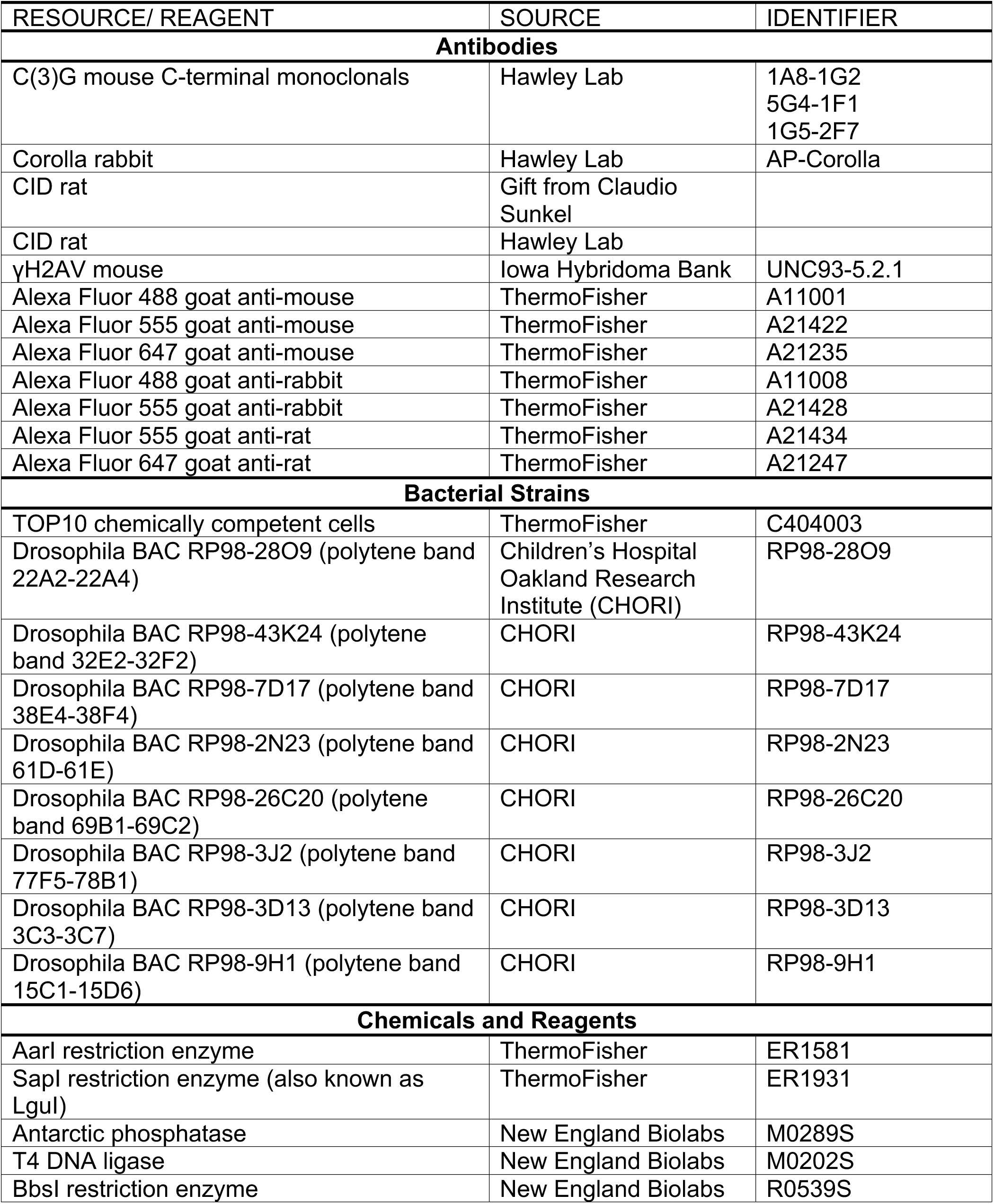
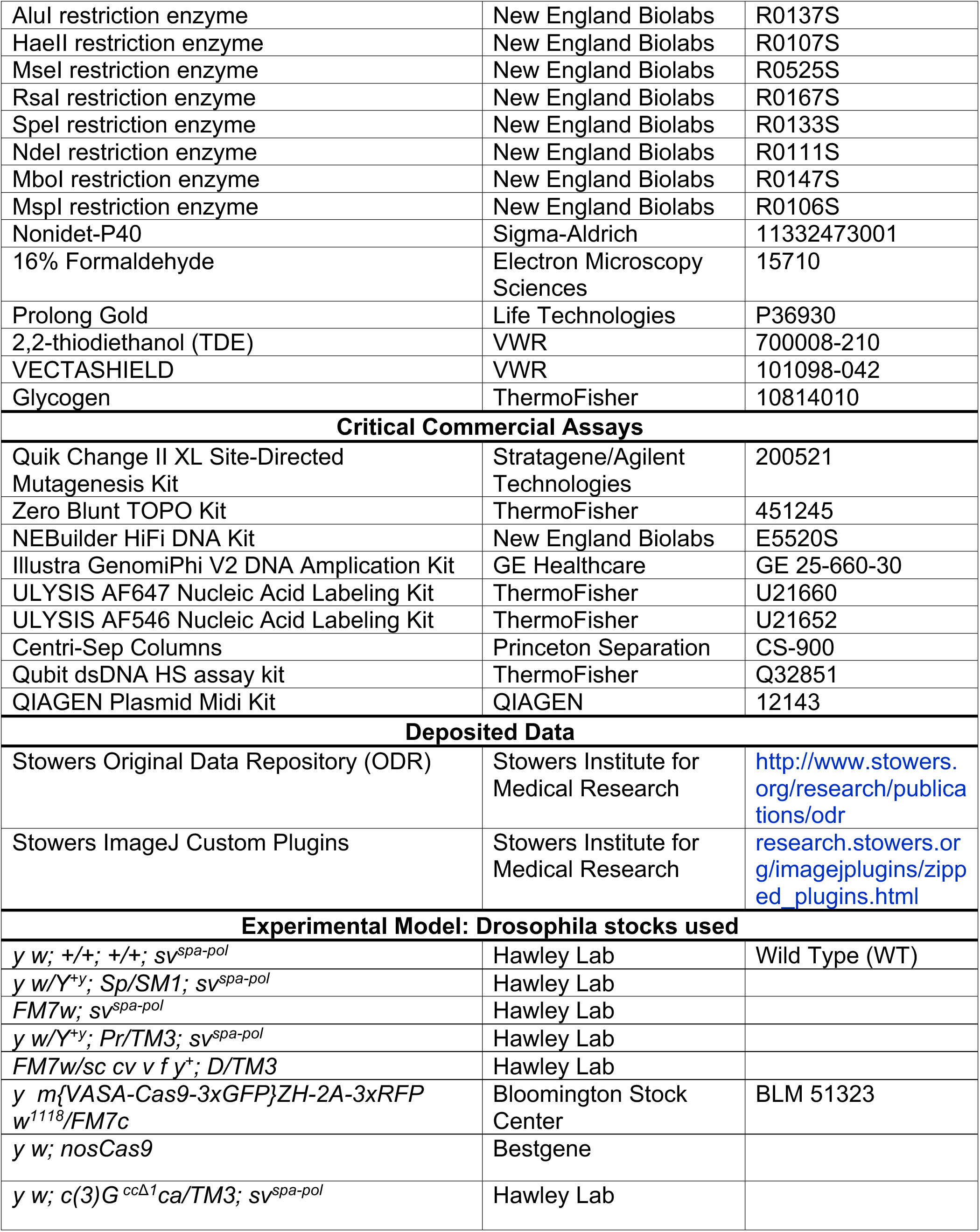
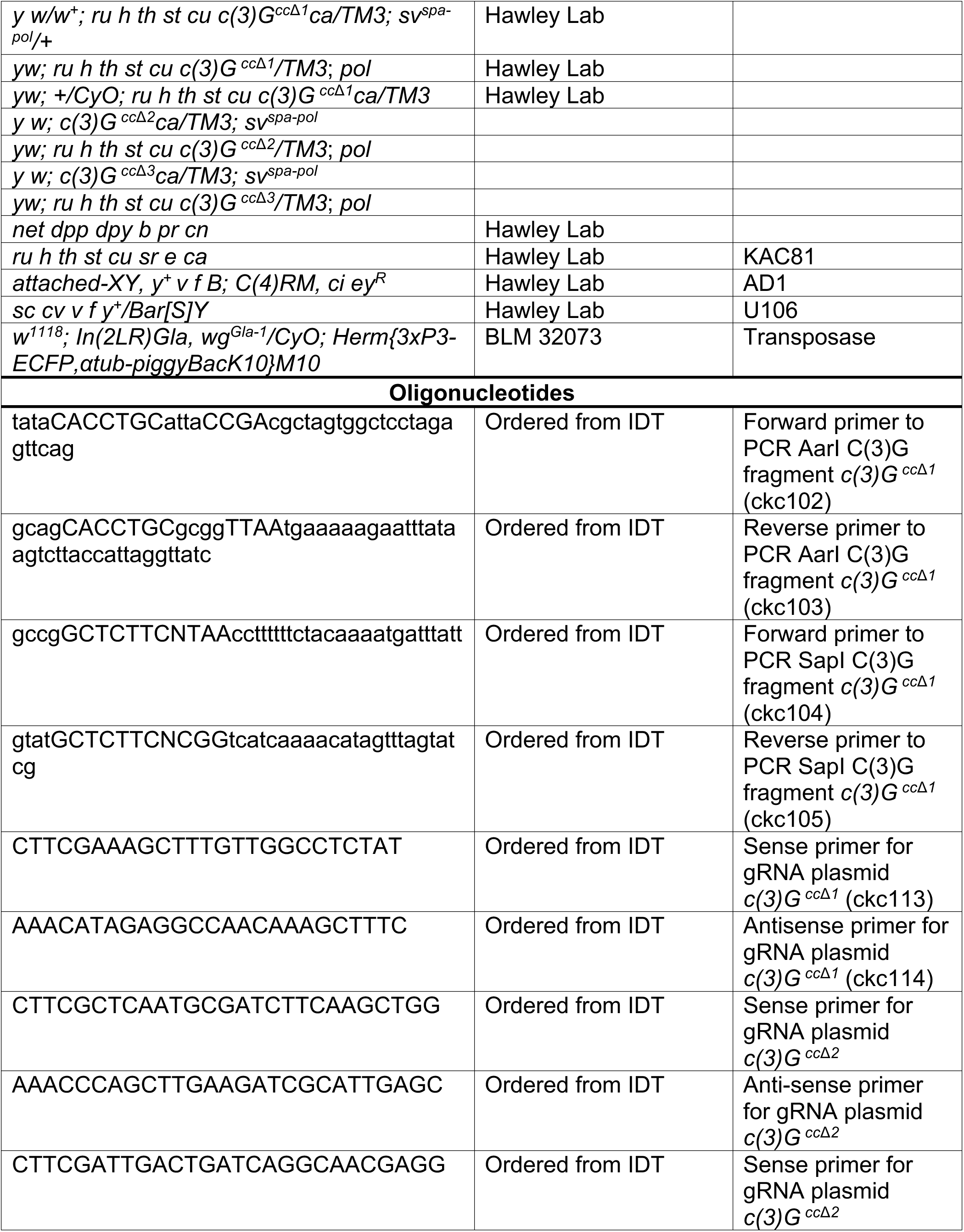
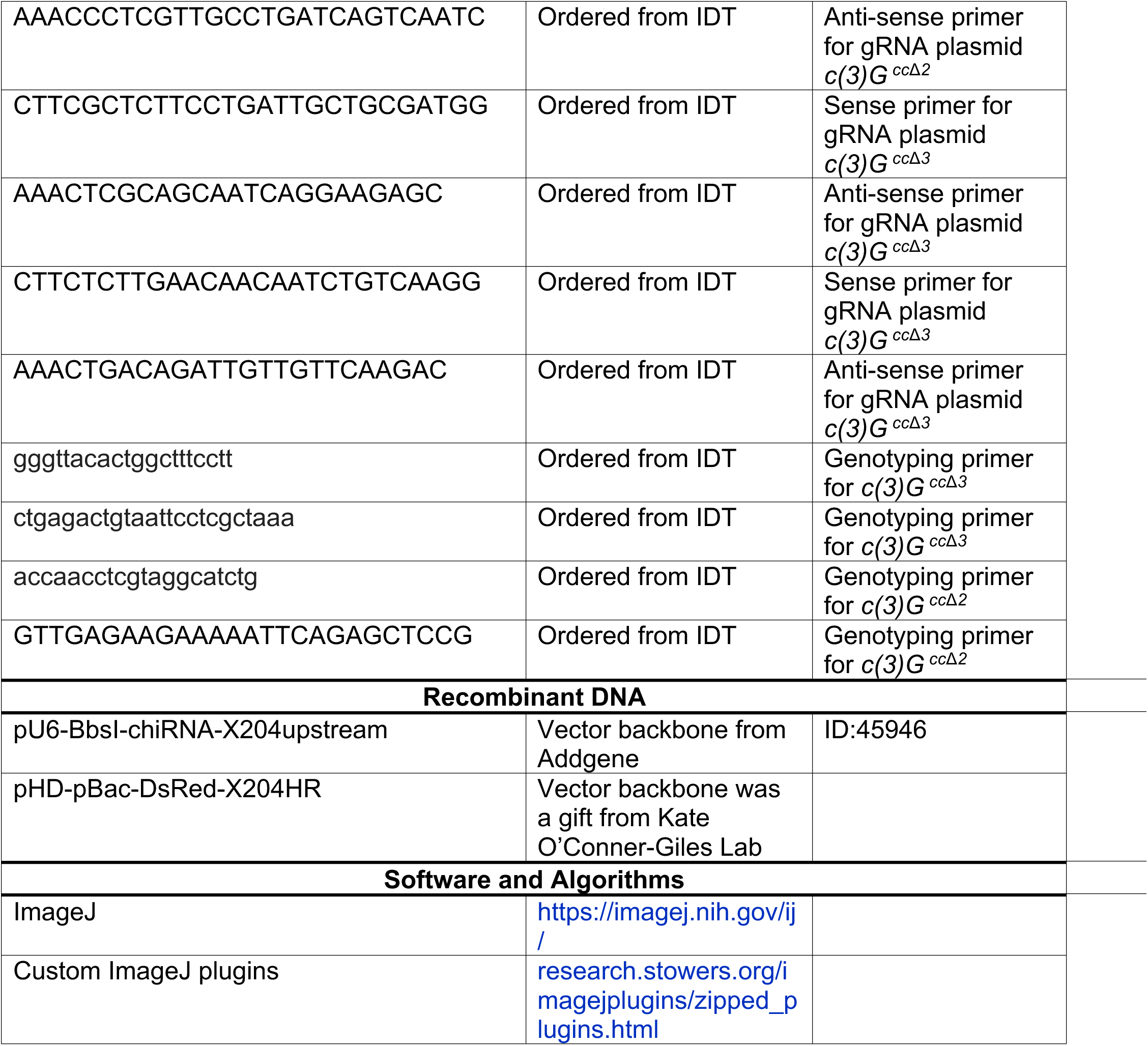
Key Resource Table.

